# Insulin resistance alters cortical inhibitory neurons and microglia to exacerbate Alzheimer’s knock-in mouse phenotypes

**DOI:** 10.1101/2025.09.05.674487

**Authors:** LaShae Nicholson, Si Jie Tang, Tejaswini Karra, Habiba Abouelatta, Stephen M. Strittmatter

## Abstract

Metabolic dysfunction contributes to the risk and progression of Alzheimer”s disease (AD) through insulin signaling, but the cellular mechanisms are not fully understood. In this study, we examined the effects of streptozotocin-induced insulin deficiency or a high-fat, high-sugar (HFHS) diet-induced insulin resistance on cognitive function in knock-in AD mouse models expressing human mutant APP and wild-type tau. Both metabolic perturbations caused hyperglycemia, but only the HFHS diet resulted in weight gain and greater learning and memory deficits. The HFHS diet exacerbation occurred without changes in amyloid-β or phospho-tau accumulation and with only subtle alterations in microglial morphology. The basis for functional deficits was explored with single-nucleus transcriptomic analysis. Prominent gene expression changes in glial cells and cerebral cortex Layer 2 inhibitory neurons correlated with the enhanced behavioral deficits. In HFHS-fed AD mice, we observed a shared *metabolic impairment in neurodegeneration* (MinD) state across multiple glial cell types. Additionally, the HFHS diet, with or without AD pathology, induced selective upregulation of the transcription factor Meis2 in cortical Layer 2 inhibitory neurons, in association with pathways involved in cell excitability. Overall, these findings suggest that HFHS-driven metabolic stress affects brain function and behavior through specific cellular programs distinct from amyloid or tau pathology, and identifies new targets that link diet-induced metabolic stress to cognitive decline in AD.

---

Dysregulation in glucose metabolism has been implicated in Alzheimer”s disease (AD) risk and progression of cognitive decline^1–3^. Insulin plays a crucial role in regulating glucose uptake and maintaining energy homeostasis across both peripheral and central systems^4–7^. Impaired insulin signaling, as the hallmark of diabetes mellitus, disrupts molecular pathways involving amyloid-β (Aβ) clearance and tau hyperphosphorylation, potentially contributing to synaptic dysfunction in AD. Metabolic stress in AD overlaps with other features of diabetes, including oxidative stress, mitochondrial dysfunction, and vascular impairment, leading to the description of AD as “type 3 diabetes”^2,8–10^. In diabetes mellitus, disrupted insulin signaling has been associated with increased Aβ accumulation in the brain. Large-scale epidemiological studies support this mechanistic link, showing that even modest elevations in blood glucose are predictive of increased AD risk and more rapid cognitive decline with aging^1^.

These observations implicate systemic metabolic dysfunction as a significant contributor to brain vulnerability in neurodegeneration. However, the cellular mechanisms linking systemic metabolic dysfunction to neurodegeneration remain poorly understood. The overlapping metabolic disruptions in AD and diabetes are particularly evident in individuals with metabolic syndrome (MetS)^11,12^, the clinical cluster of insulin resistance, obesity, dyslipidemia, and hypertension. MetS is associated with early AD onset and a fivefold higher incidence of type 2 diabetes^10,11^. MetS has been associated with systemic low-grade inflammation, vascular dysfunction, and impaired glucose regulation, features individually linked to AD^11^. Nonetheless, standard clinical measures of metabolic dysfunction, such as fasting glucose, triglyceride levels, and body mass index (BMI), are not robust predictors of cognitive outcomes. The recognition that impaired insulin signaling underlies both metabolic and neurodegenerative disorders^13–15^ has sparked interest in insulin-modulating therapies, including GLP-1 receptor agonists and intranasal insulin for AD^6,16,17^. Despite the high level of interest, the basis for systemic metabolic dysfunction altering brain function during AD progression remains uncertain.

To address this knowledge gap, we modeled metabolic perturbation *in vivo* using two well-characterized approaches to induce hyperglycemia. The administration of streptozotocin (STZ) to damage pancreatic β-cells and impair insulin production^18,19^, or the chronic consumption of a high-fat, high-sugar (HFHS) diet to induce insulin resistance^20,21^. We tested these metabolic perturbations in a disease-relevant context, using AD knock-in mice that express human mutant APP and human wild-type tau protein^22–26^. Previous studies have examined metabolic dysfunction in AD models, but most have relied on APP or tau overexpression or combined metabolic insults such as STZ plus HFHS exposure^27,28^. These approaches, along with variability in genetic context, age, and methods for inducing hyperglycemia, have produced inconsistent results and may amplify or obscure subtle effects of metabolic stress^29^. By applying distinct metabolic challenges to genetically defined AD knock-in models with pathophysiologically relevant mutations, we aimed to elucidate how metabolic dysfunction impacts behavior, glial activation, protein aggregate accumulation, and neuronal gene expression.

To evaluate metabolic, behavioral, and cellular outcomes, we combined glucose and weight monitoring, spatial memory testing, histological analyses of glial reactivity and synaptic integrity, and single-nucleus RNA sequencing (snRNA-seq) to resolve cell-type-specific transcriptional responses. Glial cell populations and cortical inhibitory neurons emerged from snRNA-seq analyses as being particularly responsive to diet-induced metabolic stress in the setting of neurodegeneration. Our findings demonstrate that while both STZ and HFHS treatments induce hyperglycemia, only the HFHS condition leads to cognitive impairments in DKI mice. This divergence suggests that metabolic stress driven by insulin resistance and obesity, hallmarks of MetS, plays a more critical role in exacerbating AD-related cognitive decline than hyperglycemia alone. Using transcriptomic profiling, we uncovered a distinct glial state associated with metabolic impairment in the AD setting, which we term the *metabolic impairment in neurodegeneration* (MinD) state. We also identified the selective upregulation of Meis2 in cortical layer 2 inhibitory neurons, induced by metabolic dysfunction alone and exacerbated by AD pathology. These results highlight how metabolic dysfunction driven by diet can modulate AD-related phenotypes independent of classical Aβ and tau pathology, offering new insight into how systemic metabolic states influence cognitive brain function.

## Results

### HFHS-Induced metabolic impairment drives cognitive decline

To investigate how impaired glucose metabolism affects AD cognitive decline, we examined reduced insulin production due to STZ-induced pancreatic β-cell loss (Extended Data Fig. 1), and chronic exposure to HFHS diet to induce insulin resistance (Fig. 1). In the STZ cohort, both STZ- and vehicle-treated wild-type (WT) and homozygous *App*^*NL-F*^*/Mapt*^*hMAPT*^ mice maintained stable body weights throughout the study (Extended Data Fig. 1a). In vehicle-treated groups, resting blood glucose remained stable over 20 weeks (Extended Data Fig. 1b). In contrast, STZ-treated mice developed fasting glucose levels >200 mg/dL within two weeks, which increased to >300 mg/dL by 20 weeks post-treatment. This models a type 1 diabetes-like phenotype, where chronic hyperglycemia results from impaired insulin production^20,21,30^. Glucose tolerance testing confirmed metabolic impairment of STZ-treated mice, which exhibited a nearly two-fold increase in blood glucose following an oral bolus glucose challenge compared to controls (Extended Data Fig. 1c). To evaluate cognitive outcomes, mice underwent Morris Water Maze (MWM) testing at 10 months (Extended Data Fig. 1d-i). STZ-treated WT and AD mice exhibited longer latencies to locate the hidden platform during initial acquisition and reversal learning (Extended Data Fig. 1d, g). However, there were no group differences compared to vehicle-treated control mice in final training sessions and probe trials (Extended Data Fig. 1e-f, h-i). All groups performed similarly in the visible platform test, confirming intact visual acuity and motor function (Fig. 1d). These results indicate that STZ-induced peripheral insulin deficiency does not exacerbate spatial learning or memory impairments in this model.

**Fig. 1.**
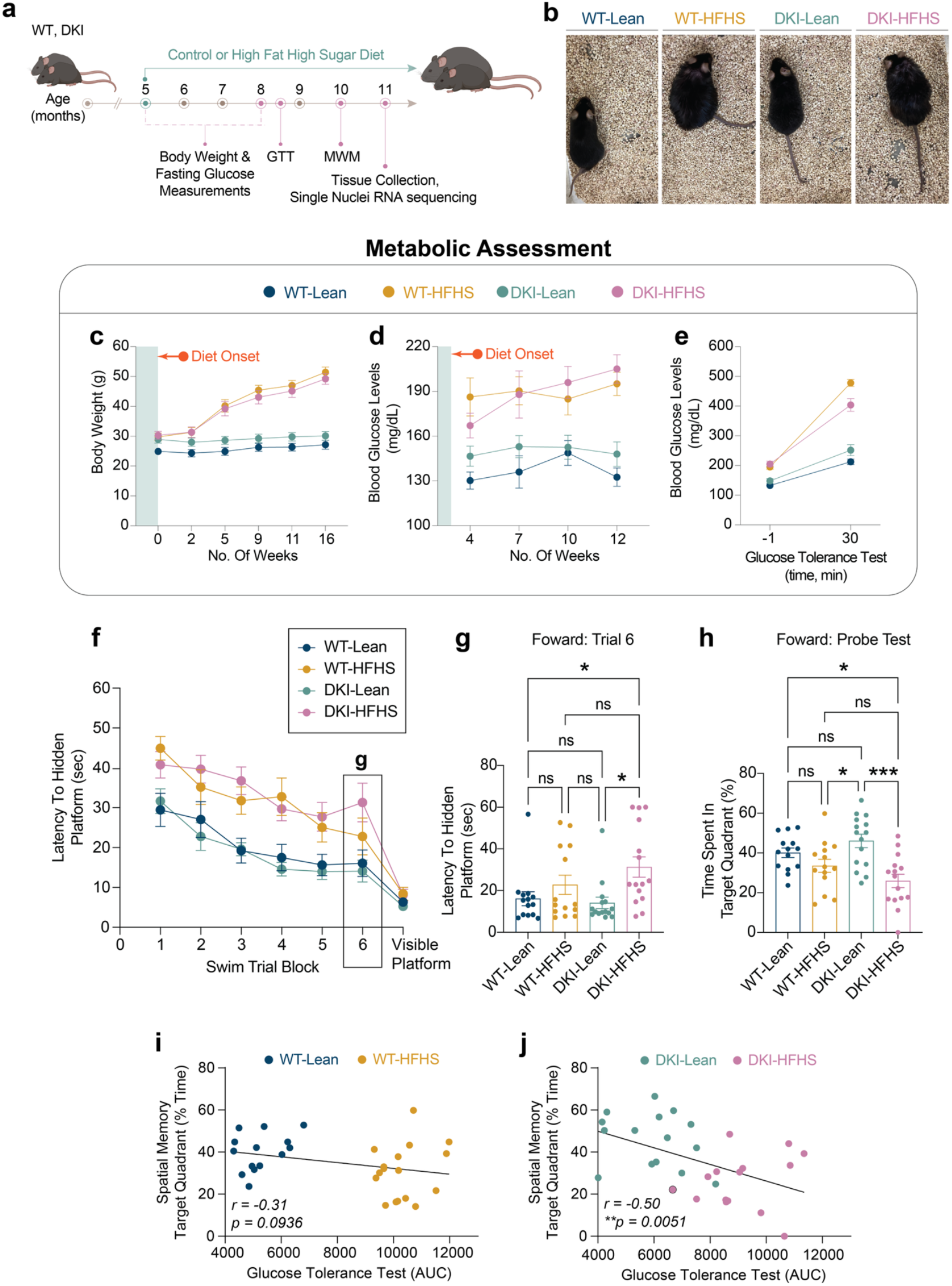
AD phenotype and HFHS diet-induced glucose intolerance synergistically impair spatial learning in DKI mice. **a-c**, Overview and metabolic assessment of the HFHS study. **a**, Schematic of experimental design showing timepoints for behavioral and metabolic assessments. **b**, Representative images of mouse body size differences between groups. **c**, Body weight **d**, blood glucose levels **e**, and glucose intolerance test 30 minutes post oral glucose bolus delivery. **f-h**, MWM behavioral testing. **f-g**, Latency time to the hidden platform during the training phase (**f**) and final training session (**g**). **h**, Fraction of time spent in the target quadrant, during acquisition probe trials. Data is presented as the mean ± SEM and analyzed by two-way ANOVA with Tukey”s multiple comparisons test, \**p<*0.0332, \*\**p<*0.0021, \*\*\**p<*0.0002, and \*\*\*\**p<*0.0001; ns: insignificant. (*n=14* for WT-Lean and WT-HFHS groups; *n=15* for DKI-Lean and DKI-HFHS groups). **i-j**, Spatial memory performance (shown in **c**) is inversely correlated with glucose intolerance in DKI (**j**) but not WT mice (**i**). Data related to Extended Data Fig. 2 and 10. MWM, Morris Water Maze (MWM); AUC, area under the curve.

We next assessed the phenotypic effects of diet-induced insulin resistance in the HFHS cohort (Fig. 1a and Extended Data Fig. 2). Unlike STZ-treated mice, both WT and homozygous *App*^*NL-G-F*^*/Mapt*^*hMAPT*^ (DKI) mice on a chronic HFHS diet exhibited significant weight gain compared to lean-diet-fed controls (Fig. 1b). Within four weeks, body weights were significantly different from those of lean-diet mice and continued to rise over 16 weeks, accompanied by elevated resting glucose levels (Fig. 1c), indicating insulin resistance-induced hyperglycemia. Glucose tolerance tests showed impaired glucose clearance in HFHS-fed mice compared to lean-fed mice (Fig. 1e), confirming the establishment of a type 2 diabetes phenotype in HFHS diet mice.

Cognitive performance in 10-month-old HFHS-fed mice was assessed using the MWM (Fig. 1f-g and Extended Data Fig. 2). During forward training, both WT-HFHS and DKI-HFHS mice had longer swim latencies to locate the hidden platform compared to lean-fed mice (Fig. 1f). However, by the final training session, only DKI-HFHS mice continued to show prolonged latencies (Fig. 1g). In the forward probe trial, DKI-HFHS mice spent significantly less time in the target quadrant than all other groups, indicating impaired memory recall (Fig. 1h). During reversal training, only DKI-HFHS mice showed delayed acquisition by the final session (Extended Data Fig. 2a-b), though no differences were observed in the reversal probe trial (Extended Data Fig. 2c). Performance in the visible platform task remained consistent across all groups (Fig. 1d). Therefore, deficits in DKI-HFHS mice reflect impaired spatial learning and memory rather than visual or motor dysfunction.

Taken together with the STZ results, the findings demonstrate that cognitive outcomes are shaped by metabolic stress driven by insulin resistance, rather than hyperglycemia being the primary factor. Both WT-HFHS and DKI-HFHS mice showed learning delays. Yet, only DKI-HFHS mice exhibited probe trial deficits, demonstrating a synergistic interaction of metabolic state and AD pathology in impairing memory function. Notably, DKI mice on a lean diet displayed intact cognitive function, in contrast to previous data for this strain on a standard diet^26,31^, further emphasizing the role of dietary intake for models of AD-related cognitive decline.

### HFHS diet alters Trem2, but not classical AD pathology

Given previous evidence of glial-mediated synapse loss in DKI mice^31^, we explored whether metabolic dysfunction modifies disease-associated microglial (DAM)^32,33^ activation state in 11-month-old mice (Fig. 2). To evaluate whether there was DAM exacerbation in DKI-HFHS mice, we examined co-immunolabeling of Iba1 and Cd68 expression in the mouse cortex (Fig. 2a-c). Cd68 immunoreactivity was elevated to a similar extent in both DKI-Lean and DKI-HFHS mice compared to their WT counterparts (*p<0*.*0001*, Fig. 2b), indicating greater microglia phagocytic activity associated with Aβ pathology. However, no significant differences were observed between DKI-Lean and DKI-HFHS mice. Iba1^+^ cell density was modestly, but significantly, increased in DKI mice relative to WT controls, independent of diet (*p*<0.0332; Fig. 2c), consistent with prior reports of gliosis in this mouse model.

**Fig. 2.**
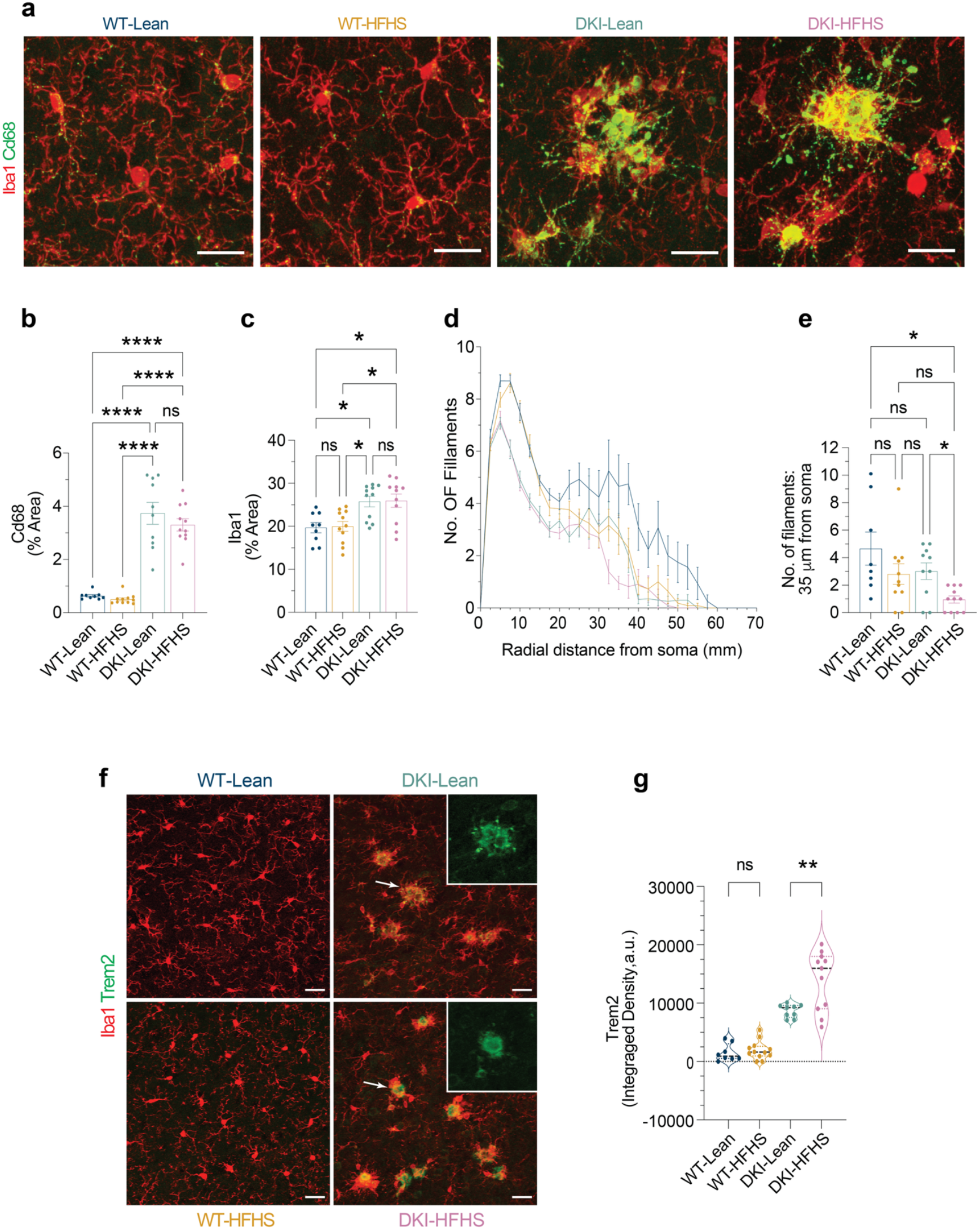
HFHS diet enhances DAM-like morphology and Trem2 proteolytic processing in DKI mice. **a**, Immunofluorescence staining showing the expression levels of Cd68 (green) in microglia (Iba1, red) within the cortex of WT and DKI mice on Lean and HFHS diet. Scale bars, 25 μm. **b-c**, Quantification of Cd68 (**b**) and microglia (**c**) percent coverage areas shown in **a**. (*n=9* for WT-Lean group; *n=11* for WT-HFHS, DKI-Lean and DKI-HFHS groups). **d**, Sholl analysis showing microglia branching complexity across treatment groups. **e**, Quantification of the number of microglia filaments at radial distances 35 μm from the soma. (Date related to Extended data Fig. 3) **d-e**, Data represents an average of 4-6 cells quantified per mouse, with each point in **e** representing a single mouse. **f**, Representative images of anti-Iba1 (red) and anti-Trem2 (green) immunostaining across treatment groups. Disease-associated Trem2 expression is elevated but exhibits an altered spatial distribution in DKI-HFHS microglia, as shown in the image insets. Scale bar, 25 µm. **g**, Quantification of TREM2 particle integrated signal density as shown in **f**. (**a**,**f**, *n=8* for WT-Lean and WT-HFHS groups; *n=11* for DKI-Lean and DKI-HFHS groups). All data presented as the mean ± SEM and analyzed by two-way ANOVA with Tukey”s multiple comparisons test, \**p<*0.0332, \*\**p<*0.0021, \*\*\**p<*0.0002, and \*\*\*\**p<*0.0001; ns: insignificant.

Although these DAM activation^33^ markers did not differ between DKI-Lean and DKI-HFHS mice, we observed subtle alterations in the distal microglial processes of WT-HFHS mice (Fig. 2a). To further evaluate whether the HFHS diet induced microglial morphological remodeling, we performed Sholl analysis on Iba1-labeled cells (Fig. 2d-e; Extended Data Fig. 3a). At 10 µm from the soma, arborization was significantly reduced in both DKI groups compared to WT (*p< 0*.*0001*; Fig. 2d, Extended Data Fig. 3a–b), consistent with disease-associated remodeling. In contrast, at 35 µm, distal microglia arborization was selectively reduced in DKI-HFHS mice (*p<0*.*0332*; Fig. 2e), with a similar but non-significant trend in WT-HFHS mice, indicating additional diet-specific effects on microglia morphology. While Sholl analysis reveals that HFHS diet consumption specifically diminishes distal branching, other global measures of microglia complexity, such as total process length, branch number, and terminal points, did not differ between DKI-Lean and DKI-HFHS groups (Extended Data Fig. 3c-e). Together, these findings suggest that while disease drives the global retraction of microglial arbors, the HFHS diet further remodels arbor geometry by promoting a more spatially confined branching pattern.

These HFHS-induced morphological changes suggested a shift in microglial activation state, potentially modifying or enhancing DAM-related physiology. We therefore examined whether the triggering receptor expressed on myeloid cells 2 (Trem2), a transmembrane receptor explicitly expressed in brain microglia and known AD risk gene regulating microglial activity^34–36^, was differentially regulated in DKI-HFHS mice. To visualize Trem2 surface expression, we used an immunolabeling strategy targeting its extracellular domain. As expected, microglia Trem2 expression was upregulated in DKI mice (Fig. 2f, see also Fig. Extended Data Fig. 5c-d). Notably, we also observed a diet-induced change in the spatial distribution of Trem2 (Fig. 2f, image insets). In DKI-Lean mice, the Trem2 signal was scattered throughout periplaque regions, whereas in DKI-HFHS mice, it was more concentrated in compact puncta, resulting in a significant increase in integrated signal intensity (*p*<0.0021; Fig. 2f,g). The remodeling of DAM morphology and shift in Trem2 localization demonstrate diet-induced modulation of microglia in the context of an AD model.

Trem2 has been implicated in the extent of both Aβ and phospho-Tau accumulation^37–39^. Using Imaris-based 3D reconstruction analyses (Extended Fig. 4A), we quantified the volume and distribution of Aβ plaques in proximity to microglia to assess potential changes in microglia-plaque interactions. Increased Aβ accumulation occurred in a DKI-dependent manner (Extended Data Fig. 4b). However, volumetric measures of Aβ were indistinguishable between DKI-Lean and DKI-HFHS mice (Extended Data Fig. 4c), and there were no diet-induced changes in microglial plaque association (Extended Data Fig. 4d). We also assessed for HFHS-diet-induced changes in neurite dystrophy and synaptic densities, since soluble Trem2 (sTREM2) has been implicated in these phenotypes as well^38^. Increases in tau phosphorylation markers, AT8 and pThr217, were observed around Aβ-plaques and Trem2 deposits in DKI mice, but with no significant difference between DKI-Lean and HFHS diet-fed mice (Extended Data Fig. 5a-d). We did observe a significant reduction in synaptic density in DKI-HFHS mice compared to WT-Lean (*p*<0.0332) and WT-HFHS (*p*<0.0021) groups, but no significant difference relative to DKI-Lean mice (Extended Data Fig. 5e,f). The absence of significant synaptic deficits in DKI-Lean mice, which aligns with their preserved spatial memory performance (Fig. 1d), differs from our previous analysis of DKI mice on a standard laboratory diet^26,31^ as well as the HFHS diet, and suggests that dietary restriction may protect vulnerable synapses from disease pathology. Despite the documented shifts in microglial and synaptic metrics, overall Aβ deposition and tau phosphorylation remained unaffected by HFHS consumption in DKI mice.

### HFHS induces a glial metabolic impairment state in DKI mice

To further investigate glial regulation by metabolic state, we performed single-nucleus RNA sequencing (snRNA-seq) to evaluate transcriptomic profiles in depth. Pooled cortical and hippocampal tissues from individual mice (*n=4* per group) were processed for nuclei enrichment and isolation. Samples were sequenced, integrated, and processed for batch-corrected dimensional reduction in UMAP space, followed by Leiden clustering (Extended Data Fig. 6a). Clusters were annotated with major cell type labels based on cluster-specific marker genes, which were identified by differential expression analysis (Extended Data Fig. 6b). Annotations were cross-validated using the MapMyCells (RRID: SCR_024672) tool from the Allen Brain Cell Atlas (Supplemental Table 1). Post-processing, the dataset yielded a mean sequencing depth of 450 million reads per group, with an average of 30,000 reads and approximately 1,500 unique molecular identifiers (UMIs) per nucleus (Extended Data Fig. 6d-f). Several populations of excitatory (ExN) and inhibitory (InN) neurons were identified, along with various glial and non-glial blood-brain barrier cells, with each cell type represented across the groups (Extended Data Fig. 6c). While the proportions of neuronal nuclei remained consistent across conditions, the frequencies of glial populations varied between groups. The number of oligodendrocyte nuclei increased with diet, disease, and their combination, while DKI mice showed elevated microglia and reduced astrocyte frequencies, in line with prior findings^26,31^. In UMAP space, a subset of inhibitory neurons appeared exclusively in HFHS-fed mice (open arrows), indicating a diet-specific response that is examined below. First, we focused on microglia to identify transcriptional changes that may underlie the observed Trem2 alterations, despite the equal number of microglial nuclei in the DKI-Lean and DKI-HFHS groups (Extended Data Fig. C).

Microglial nuclei were subsetted and reclustered for downstream analysis, revealing the selective enrichment of distinct microglial subpopulations across experimental groups (Fig. 3a; Extended Data Fig. 7a,b). Leiden clustering yielded 11 microglial subclusters, as shown in UMAP space (Fig. 3a). Clusters MG8, MG9, MG10, and MG11 predominantly consisted of homeostatic microglia, while clusters MG4, MG5, and MG7 were enriched in DKI mice, expressing marker genes consistent with previously described DAM profiles (Extended Data Fig. 7b,c). Notably, cluster MG3 was selectively enriched in DKI-HFHS mice (arrows in Fig. 3a,c), representing a unique transcriptional state associated with metabolic stress in the context of neurodegeneration, which we refer to here as a *<u>m</u>etabolic <u>i</u>mpairment in <u>n</u>euro<u>d</u>egeneration* (MinD) state. Differential expression analysis (complete list in Supplementary Table 3) identified *Dlg2, Kcnip4, Lsamp, Ptprd, Nrg3*, and *Clec4f* as the top marker genes selectively enriched and upregulated in MG3 relative to all other microglial clusters (Fig. 3c). We next cross-referenced gene expression within the MG3 cluster against a curated list of microglial state markers^33^, which were stratified by WT and DKI groups based on disease-associated activation (Extended Data Fig. d). In contrast, MG3-enriched MinD genes were upregulated explicitly in DKI-HFHS microglia, indicating a metabolically impaired state driven by the combined effects of disease and diet (Extended Data Fig. 7c,d).

**Fig. 3.**
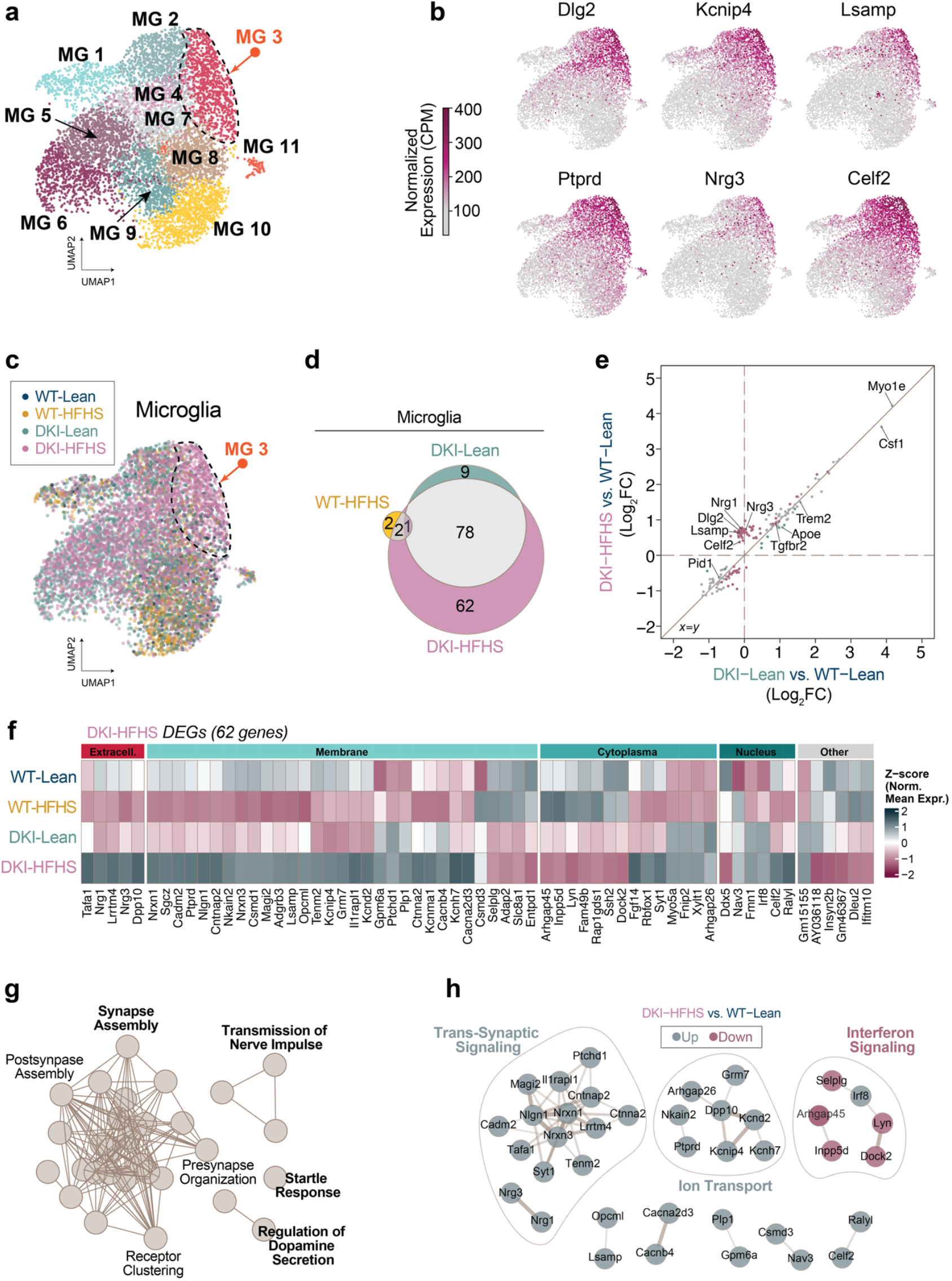
HFHS induces a metabolic impairment transcriptional program atop of DAM gene signature. **a-h**, snRNA-seq analysis of microglial nuclei populations isolated from 11-month-old WT and DKI mice under lean or HFHS conditions (*n=4* mice per group). **a**, Subclustering of microglial nuclei shows distinct transcriptomic subsets. Cluster MG3 (orange arrow) exhibits predominant enrichment of DKI-HFHS microglia. In contrast, other clusters display overlapping contributions from both DKI and WT groups across dietary conditions (related to Extended Data Fig. 6). **b**, Nuclei transcript levels of the top six marker genes for the MG3 cluster. **c**, Overlay of group nuclei expression profiles in UMAP space. **d**, Venn diagram illustrating the overlap of shared DEGs from WT-Lean in WT-HFHS, DKI-Lean, and DKI-HFHS groups. **e**, Correlation plot comparing Log_2_ fold-changes of DEGs in DKI-Lean and DKI-HFHS microglia relative to WT-Lean microglia. **f**, Cellular localization and group expression patterns of the 62 DKI-HFHS-specific DEGs identified in **d. g**, Pathway enrichment analysis of 62 DKI-HFHS-specific DEGs depicted in **d**-**f. h**, PPI network of genes depicted in **d-f** with MCL clustering. Nodes represent individual proteins, while edges represent the interaction between proteins. Color-coded nodes indicate upregulated (green) and downregulated (magenta) genes. Edge thickness represents the relative protein interaction strength. Theme labels represent the main GO Biological Process PPI of the respective subcluster. See Supplemental Table 3 for the complete list of DEGs with corresponding Log_2_ fold-change values, pathway enrichment, and PPI analysis outputs. See also Extended Data Fig. 5–7. UMAP, Uniform Manifold Approximation and Projection; CPM, counts per million; DEGs, differentially expressed genes; PPI, protein-protein interaction; MCL, Markov Cluster algorithm.

**Fig. 4.**
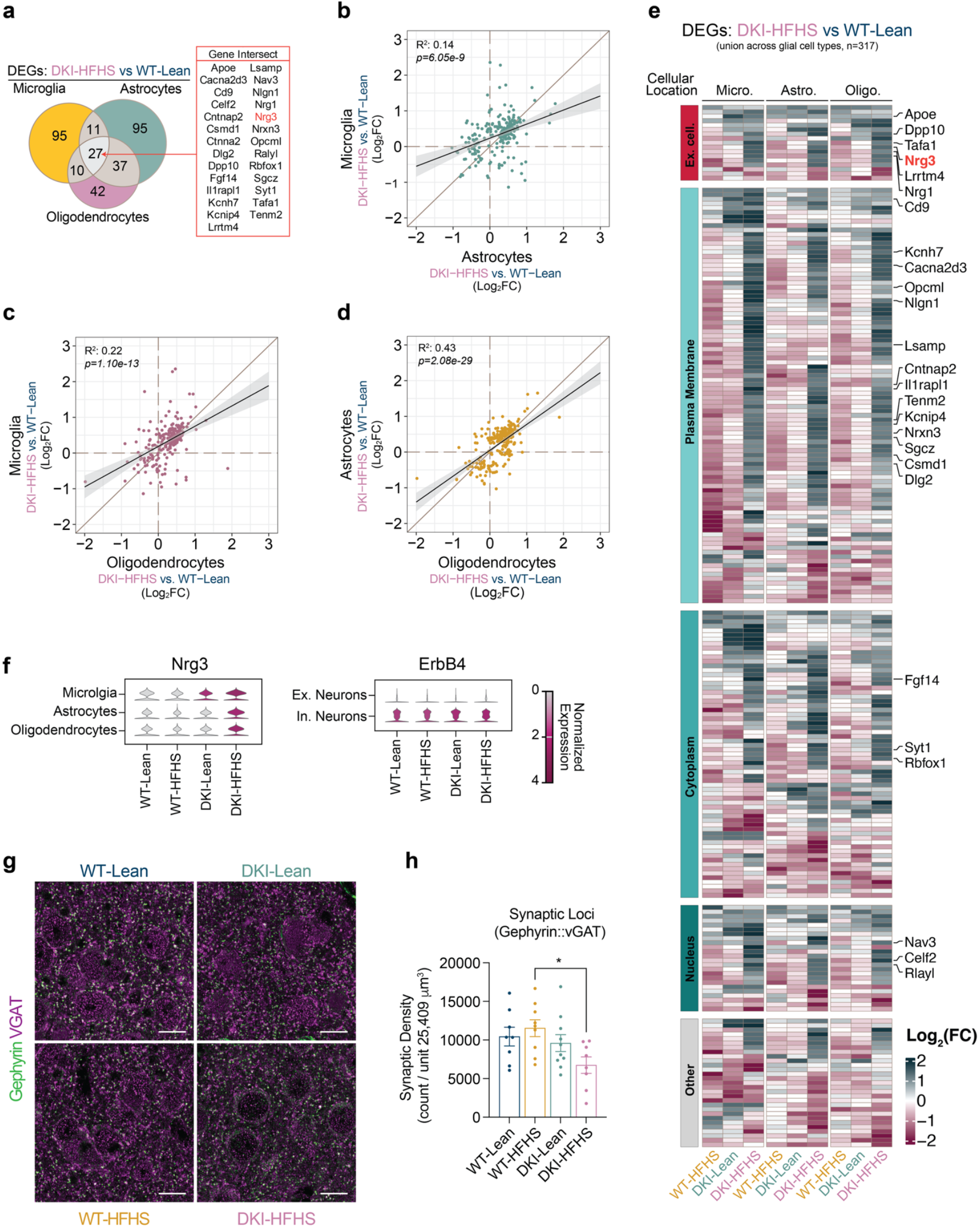
Loss of inhibitory synapses in DKI-HFHS mice corresponds with pan-glial MinD state. **a-e**, Shared DEGs across microglia, astrocytes, and oligodendrocytes show positive intercellular correlations and reflect a common MinD state transcriptional program upregulated in DKI-HFHS mice. **a**, Venn diagram of shared and unique DEGs across microglia, astrocytes, and oligodendrocytes in the DKI-HFHS group, with DEGs common to all glial types listed. **b-d**, Pairwise correlation plots comparing Log_2_ fold-changes of shared DEGs from **a** for microglia vs astrocytes (**b**), microglia vs oligodendrocytes (**c**), and astrocytes vs oligodendrocytes (**d**), in the DKI-HFHS vs WT-Lean comparison, each showing a positive correlation. **e**, Union of DEGs identified in (**a-d**) organized by protein cellular location, highlighting a glial-specific transcriptional program linked to metabolic impairment in DKI-HFHS mice. Data Related to Fig. 3 and Extended Data Fig. 7. See Supplemental Tables 3 and 5 for the complete DEG list. **f**, Stacked-violin plots showing normalized transcriptomic levels of Nrg3 in glial and ErBB4 in neuronal nuclei populations. **g**,**h**, Representative immunostaining (**g**) and corresponding colocalization quantification of Gephyrin with vGAT-positive synapses (**h**). (*n=9* for WT-Lean group; *n=11* for WT-HFHS, DKI-Lean, and DKI-HFHS groups.) For (**h**) data presented as the mean ± SEM and analyzed by two-way ANOVA with Tukey”s multiple comparisons test, \**p<*0.0332, \*\**p<*0.0021, \*\*\**p<*0.0002, and \*\*\*\**p<*0.0001; ns: insignificant. MinD, *metabolic impairment in neurodegeneration*; DEG, differentially expressed genes; micro, microglia; astro, astrocytes; oligo, oligodendrocytes.

To avoid biases introduced by differences in nuclei counts within subcluster MG3, differential expression analyses were conducted on the total microglial population across groups. We performed DEG analysis comparing each experimental group to WT-Lean, applying a significance threshold of adjusted *p*<0.05 and log_2_ fold-change greater than ±0.25. A Venn diagram of significant DEGs (Fig. 3d, Supplemental Table 3) revealed 78 genes shared between DKI-HFHS and DKI-Lean groups, while 62 genes were uniquely altered in DKI-HFHS microglia. To visualize correlative gene expression patterns, we then plotted the log_2_ fold-changes for DKI-HFHS and DKI-Lean microglia relative to WT-Lean (Fig. 3e). This revealed a strong positive correlation in disease-associated gene expression, with overlapping (gray), DKI-Lean–specific (green), and DKI-HFHS–specific (magenta) genes distributed along the identity line. Notably, a distinct cluster of magenta points—representing nearly half of the DKI-HFHS–specific DEGs—deviated from this trend, suggesting additional transcriptional changes driven by the compound effects of diet and disease.

To better understand the biological context of genes specifically altered in DKI-HFHS microglia, we organized the 62 DEGs in a heatmap according to their predicted cellular localization and visualized their relative expression levels across each group (Fig. 3f). Of the specific DKI-HFHS genes clustered in the correlation plot (Fig. 3e), a substantial proportion encoded plasma membrane–associated or secreted proteins (Fig. 3f). Pathway enrichment analysis (Fig. 3g) highlighted gene sets involved in synaptic assembly, presynaptic organization, and synaptic receptor clustering. A protein–protein interaction network analysis (Fig. 3h, Supplemental Table 3) identified dense connectivity among genes involved in trans-synaptic signaling and ion transport. In contrast, a subset of genes associated with the interferon signaling pathway was downregulated, except for Irf8, a transcription factor implicated in microglial immune activation^40,41^.

Pan-glial transcriptomic remodeling was evident in the DKI-HFHS mice, extending beyond microglia to astrocytes and oligodendrocytes (Fig. 4a–h, Extended Data Fig. 8 and 9). Significantly greater GFAP-positive astrocyte signal was detected in the cerebral cortex of both DKI groups (Extended Data Fig. 8a,b), coinciding with significant disease-associated activation compared to WT controls (*p<*0.0001). However, consistent with microglia, astrocytes (Extended Data Fig. 8c) and oligodendrocytes (Extended Data Fig. 9a,b) in DKI-HFHS mice displayed transcriptomic profiles resembling a MinD-like state (Fig. 3, Extended Data Fig. 7). In UMAP space, distinct sets of genes were uniquely enriched from DKI-HFHS mice, further strengthening the evidence that astrocytes and oligodendrocytes adopt MinD-like transcriptional programs (Extended Data Fig. 8g, 9e). In astrocytes and oligodendrocytes, log_2_ fold-change comparisons relative to WT-Lean revealed discrete separation between DKI-Lean and DKI-HFHS groups. Notably, a subset of DEGs was uniquely detected in the DKI-HFHS mice, reflecting a unique transcriptional program not shared in the respective glial cell populations in DKI-Lean mice (Extended Data Fig. 8f, 9d). Differential expression analysis confirmed that DKI-HFHS mice exhibited the most significant number of transcriptional alterations across astrocytes and oligodendrocytes (Extended Data Fig. 8d, 9c). Although many changes were unique to individual populations, 27 DEGs were shared across all three glial types (Fig. 4a). Correlation analyses demonstrated strong positive associations in DEG expression between microglia, astrocytes, and oligodendrocytes (Fig. 4b-d). These overlapping genes, organized by predicted cellular localization (Fig. 4e), define a shared MinD transcriptional program in response to combined metabolic and neurodegenerative stress.

Among the MinD state genes with selective upregulation in DKI-HFHS microglia, neuregulin 3 (Nrg3) consistently emerges as a gene of interest (Fig. 3b, e, f, h). Nrg3, along with the upregulated Nrg1, belongs to the neuregulin family of secreted ligands known to modulate synaptic function through ErbB receptors^42^. Nrg3 transcripts were also upregulated in astrocytes and in oligodendrocytes of the DKI-HFHS mice (Fig. 4a,f,e; Extended Data Fig. 8e; Extended Data Fig. 9e). While Nrg1 can signal through ErbB2, 3, and 4^42–44^, Nrg3 primarily binds to ErbB4, a receptor selectively enriched at inhibitory synapses (Fig. 4f)^42,44–46^. Therefore, we assessed inhibitory synapse density using gephyrin–vGAT co-localization and observed reduced inhibitory synapse loci in DKI-HFHS mice (Fig. 4g,h, and Extended Data Fig. 5e,f). These data define one molecular pathway specifically engaged by the combination of AD pathology and insulin resistance, leading to altered neuroglial interactions.

### HFHS triggers metabolic impairment program in Meis2+ L2 neurons

We considered whether neuronal gene expression varied in parallel with these glial changes. Subclustering of inhibitory neurons revealed seven transcriptionally distinct groups, with a selective enrichment of nuclei from HFHS-fed mice in cluster InN3 (arrows, Fig. 5a,b). DEG analysis showed a high number of transcriptional changes in InN3 neurons in both WT-HFHS and DKI-HFHS mice, but not in DKI-Lean mice, compared to WT-Lean (Fig. 5c). The majority of DEGs were shared between WT-HFHS and DKI-HFHS (Fig. 5d), indicating a common transcriptional response to HFHS exposure (Fig. 5e). Pathway enrichment of the combined DEGs revealed densely interconnected nodes associated with synaptic organization, vesicle release, and trans-synaptic signaling, with the most prominent gene ontology terms linked to the regulation of membrane potential (Fig. 5f, Supplemental Table 5). These signatures point to altered cellular excitability as a defining feature of InN3 neurons under metabolic stress. Cell type classification identified InN3 nuclei as Meis2^+^ Layer (L) 2/3 inhibitory neurons (Fig. 6a-b,d-e, and Supplemental Table 1). Meis2 transcripts, like the other top five marker genes *Pbx1, Auts2, Kcnip2*, and *Pbx3*, have strongly elevated transcript levels in InN3 nuclei. *In vivo, Meis2* expression was also elevated in L2/3 cortical neurons in response to HFHS, with the most substantial increase observed in DKI-HFHS mice. As a transcription factor linked to pancreatic glucose dysregulation^47,48^, increased *Meis2* expression in cortical neurons suggests a parallel metabolic impairment program for inhibitory neurons in the brain. These findings also highlight L2/3 Meis2^+^ neurons as selectively vulnerable to HFHS-driven metabolic stress.

**Fig. 5.**
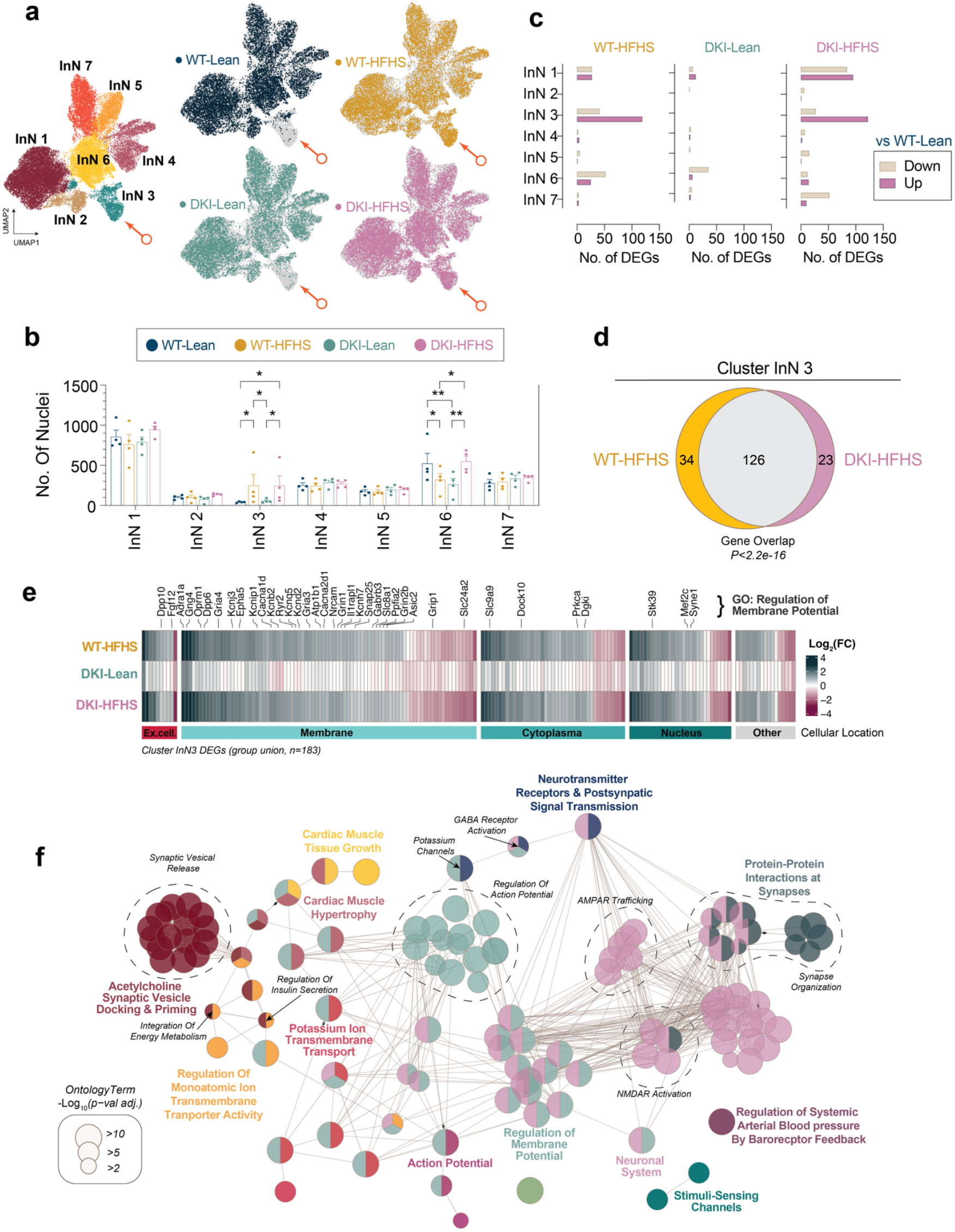
HFHS diet induces a metabolic stress transcriptional program in specific inhibitory neurons. **a**, UMAPs showing subclustering of InN nuclei (left) and group expression profiles (right). Orange arrows point to subcluster InN 3, which exhibits selective nuclei enrichment in HFHS diet groups. **c**, DEG counts for InN subclusters of each respective group vs WT-Lean. **d**, Nuclei frequencies of subclusters shown in **a. d**, Venn diagram illustrating the overlap of shared DEGs between WT-HFHS and DKI-HFHS groups shown in **b**. Significance *P-*value for gene overlap was computed using Fisher”s Exact Test. **e**, Heatmap of the Log_2_ fold-changes of the union of all DEGs depicted in **b**,**d** organized by each gene”s protein cellular location. Highlighted Log_2_ fold-changes are of all the genes represented by the *“Regulation of Membrane Potential”* node shown in **f. f**, Pathway enrichment analysis of nuclei subcluster InN 3. The Enrichment Map represents clustered nodes of GO pathways, with edges connecting pathways shared by many genes. Theme labels represent the primary *GO Biological Process* term associated with each node subcluster, with colors coordinated to enhance visualization. Node sizes correspond to the −Log_2_ transformed adjusted *p*-value for each GO term gene set enrichment. See Supplemental Table 5 for a complete list of DEGs and pathway enrichment analysis outputs. UMAP, Uniform Manifold Approximation and Projection; InN, inhibitory neurons; vs, versus. GO, gene ontology.

**Fig. 6.**
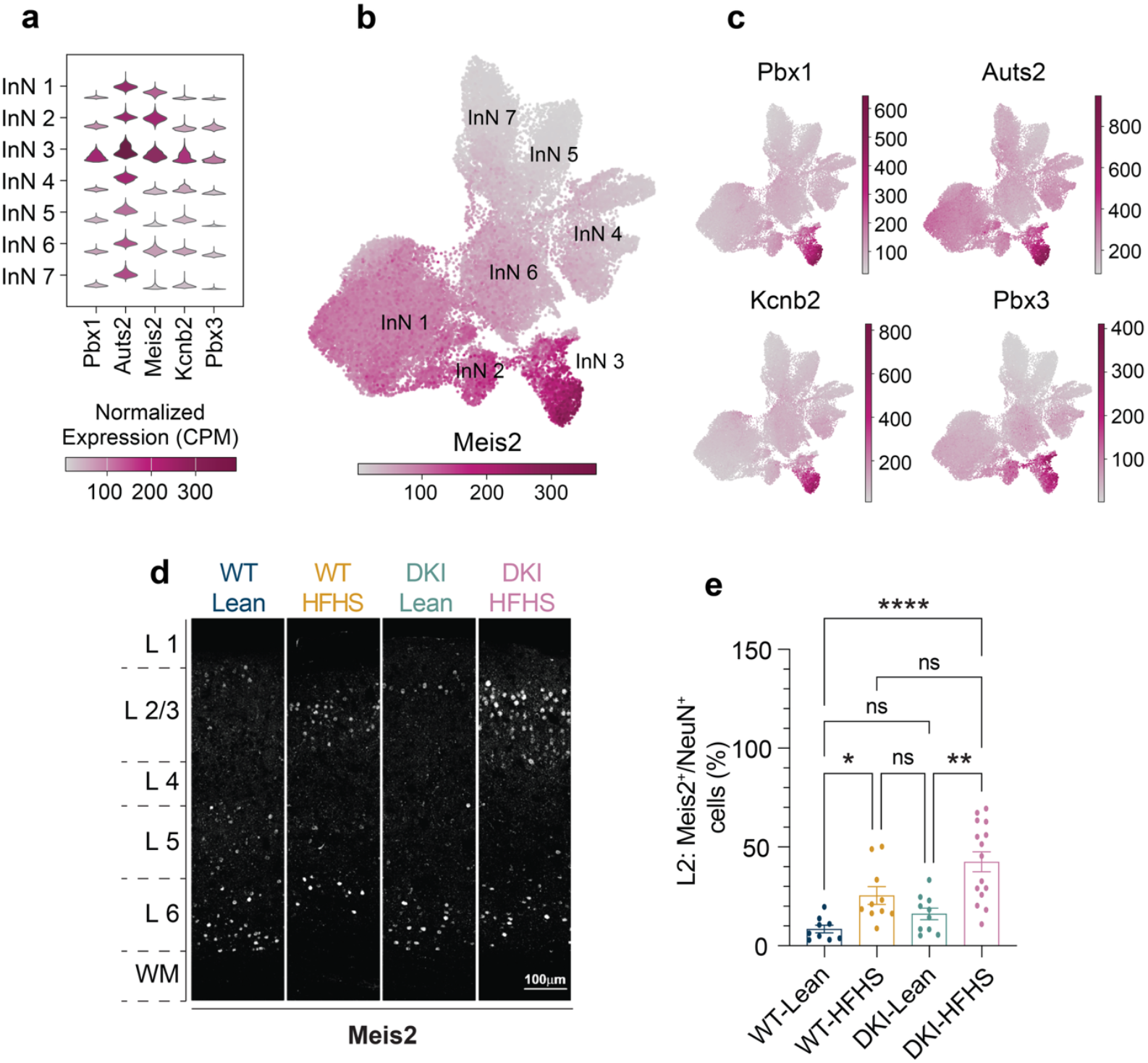
HFHS diet induces Meis2 upregulation in L2 inhibitory neurons with amplified levels in DKI mice a-c, Top marker genes for subcluster InN 3, identified as cortical L2 Meis2^+^ InNs. See Supplemental Table 5 for the full list of cluster marker genes. Data related to Fig. 9. **a**, Stacked-violin plot depicting normalized transcript levels of the top five InN 3 cluster marker genes, displayed across all InN subclusters. **b**,**c** The transcript levels of the top genes shown in **a**, displayed in UMAP space. Scale bars, normalized CPM. **d**, Representative images demonstrating Meis2 expression levels across cortical layers. There is a diet-induced increase of Meis2 within cortical L2, which is increased in DKI-HFHS mice. **e**, Quantification of the percentage of Meis2^+^ cells as a proportion of the total number of NeuN+ cells located in cortical L2 as represented in **d**. (*n=9* for WT-Lean group; *n=11* for WT-HFHS, DKI-Lean and DKI-HFHS groups.) Data presented as the mean ± SEM and analyzed by two-way ANOVA with Tukey”s multiple comparisons test, \**p<*0.0332, \*\**p<*0.0021, \*\*\**p<*0.0002, and \*\*\*\**p<*0.0001; ns: insignificant. CPM, counts per million; InN, inhibitory neurons; L2, layer 2; WM, white matter.

Recently, reports indicated that the HFHS diet promotes insulin resistance by thickening perineuronal nets (PNNs) around hypothalamic AgRP neurons, thereby limiting their response to peripheral signals, such as insulin^49^. Since PNNs are also known to ensheathe Pvalb^+^ interneurons in the cortex^50,51^, we investigated whether HFHS-associated changes in Meis2^+^ inhibitory neurons align with cortical PNN remodeling. Utilizing Wisteria floribunda agglutinin (WFA) labeling combined with Pvalb immunostaining, we found that PNNs were predominantly localized to cortical L4 (Extended Data Fig. 10a). A trend toward increased PNN density was observed in response to both diet and disease (Extended Data Fig. 10c); however, this increase was restricted to deeper cortical layers, with no PNN expansion detected in L2/3 (Extended Data Fig. 10a). Notably, Meis2^+^ neurons in L2/3 did not co-localize with Pvalb and Pvalb^+^ cell numbers remained unchanged across groups (Extended Data Fig. 10b-d). These findings suggest that the transcriptional changes observed in Meis2^+^ neurons are unlikely to result from PNN-associated insulin resistance, distinguishing them from mechanisms described in hypothalamic AgRP neurons^49^.

### ExNs have distinct yet convergent transcriptomics patterns of synaptic dysregulation

We next investigated whether excitatory neurons exhibit a transcriptomic signature of metabolic impairment in response to HFHS. Thirteen excitatory subclusters were identified (Fig. 7a), with ExN9 and ExN10 showing the highest number of DEGs following HFHS exposure (Fig. 7b). A smaller set of disease-associated DEGs was also detected in DKI-Lean neuronal nuclei. Cell type classification (Supplementary Table S) identified these clusters as L2-L4 excitatory neurons, with ExN9 enriched for L3/4 markers and ExN10 for L2/3 markers (Fig. 7c). Given the sensitivity to metabolic impairments observed in Meis2^+^ L2 inhibitory neurons, we focused our analysis on cluster ExN10, representing L2/3 excitatory neurons. DEG analysis revealed distinct, group-specific gene sets with minimal overlap (Fig. 7d, Supplemental Table 6). However, the resulting expression signatures were strikingly similar across conditions, indicating transcriptional convergence despite divergent gene identities (Fig. 7e). Merged pathway analysis with hierarchical clustering confirmed this pattern. While some functional nodes were specific to WT-HFHS (yellow) or DKI-HFHS (magenta) DEGs, the majority reflected shared contributions (gray) (Fig. 7f, see Supplemental Table 6 for all pathway enrichment outputs). Like the pathway enrichment analysis of Meis2^+^ L2 inhibitory neurons (Fig. 5f), the most highly connected pathways were associated with synapse organization and trans-synaptic signaling, driven by genes localized to synaptic membranes (Fig. 7f,g). Additionally, a node cluster of separate axon-specific pathways was uniquely enriched in WT-HFHS (Fig. 7f). These findings suggest that excitatory neurons exhibit a transcriptomic signature of metabolic impairment that converges on altered synaptic function through partially overlapping yet mechanistically distinct programs.

**Fig. 7.**
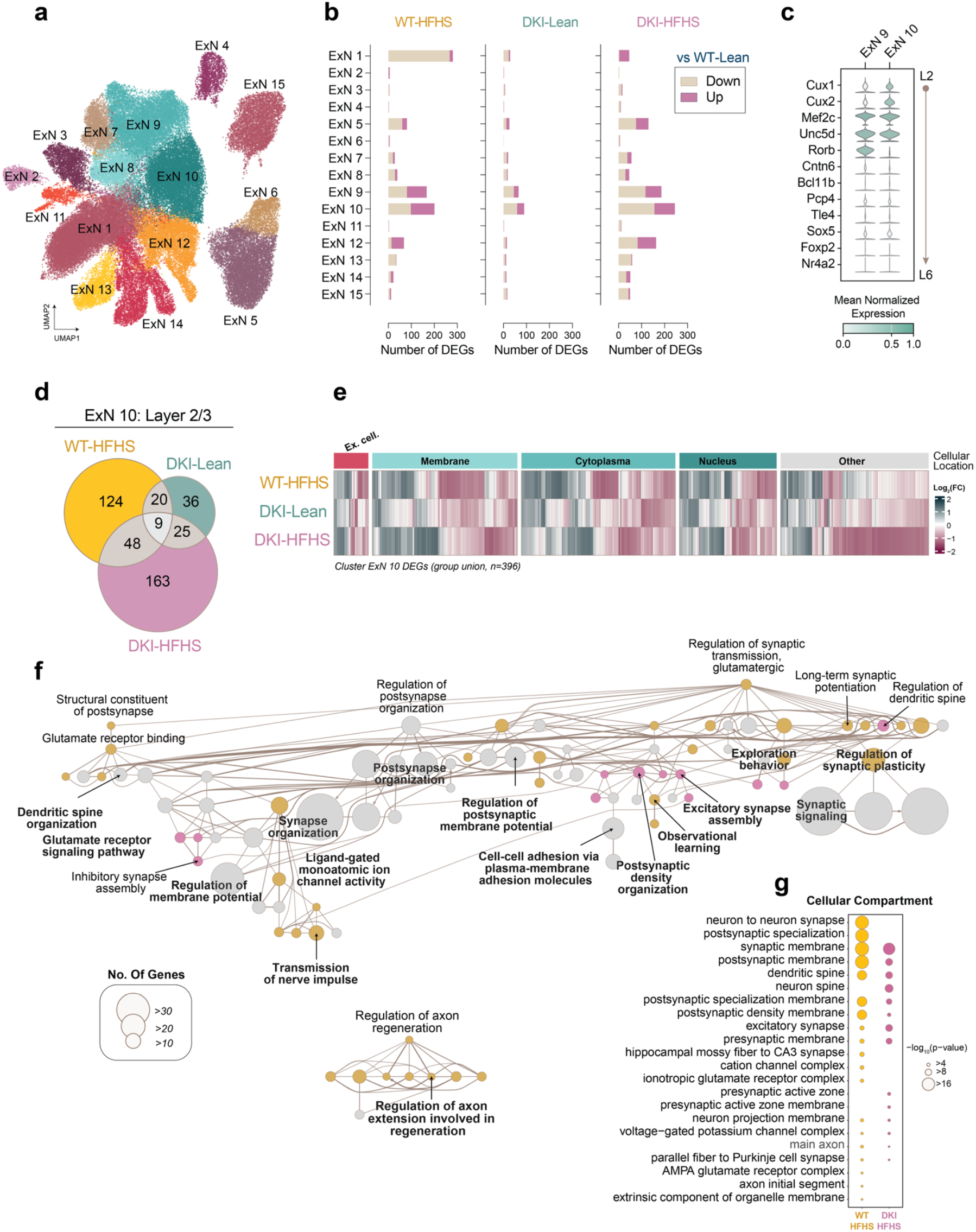
Layer 2-4 ExNs show distinct disease- and diet-specific transcriptional changes with convergent synaptic pathway dysregulation. **a**, UMAP of subclustered ExN nuclei populations. **b**, DEGs counts per ExN subcluster for each group compared to WT-Lean. ExN 9 and ExN 10 nuclei subpopulations exhibit genetic dysregulation in response to disease, diet, and the combination of the two. (See Supplemental Table 6 for full DEG list) **c**, Stacked-violin plots showing cortical layer markers corresponding to L2-L6. Marker genes for L2 and L3 are enriched in nuclei cluster ExN 10, while L3 and L4 marker genes are enriched in nuclei cluster ExN 9. **d**, Venn diagram showing DEGs overlap between groups as compared to WT-Lean mice for nuclei cluster ExN 10. **e**, Heatmap of the Log_2_ fold-changes of the union of all cluster ExN 10 DEGs depicted in **b**,**d**, organized by each gene”s protein cellular location. **f**, Merged pathway enrichment map for WT-HFHS and DKI-HFHS dysregulated genes, organized using a tree layout where nodes are arranged hierarchically to emphasize parent-child relationships between pathways. Nodes are colored by gene set contribution (yellow, WT-HFHS; magenta, DKI-HFHS; grey, equal contributions), and node size reflects the number of genes. **g**, Dot plot of GO Cellular Component terms resulting from the enrichment analysis for ExN 10 cluster DEGs for WT-HFHS and DKI-HFHS nuclei samples. Dot size indicates the *p-*value of significance of each term. ExN, excitatory neurons; DEGs, differentially expressed genes; L, layers.

## Discussion

This study identifies two key cellular responses to metabolic dysfunction in the context of AD: a shared glial MinD transcriptional program and the selective upregulation of Meis2 in cortical L2 inhibitory neurons. These cellular and molecular findings provide a basis to understand the capacity of brain insulin resistance to exacerbate AD-related neurological dysfunction. The use of double-knock-in AD mouse models, chronic diet-induced metabolic syndrome, and single-nucleus transcriptomic profiling of brain tissue strengthens the translational relevance of our observations. Together, these data underscore the importance of considering metabolic dysfunction when evaluating age-dependent AD pathophysiology.

### Metabolic syndrome as a disease modifier of Alzheimer”s progression

Metabolic dysfunction is increasingly recognized as a critical modifier of AD risk and progression^1,2,8,9,14,52^, yet the underlying mechanisms have been ill-defined. We examined how metabolic stress influences AD-related phenotypes in knock-in mice in response to both STZ-induced pancreatic β-cell loss and diet-induced insulin resistance. While both perturbations induced hyperglycemia, only the HFHS-fed DKI mice exhibited spatial learning deficits, accompanied by weight gain and insulin resistance, consistent with the clinical presentation of MetS. In contrast, STZ-treated mice exhibited persistent hyperglycemia without cognitive impairment. These findings indicate that the acceleration of AD-related cognitive decline is not driven solely by elevated glucose but by the broader physiological state of diet-induced chronic metabolic stress in response to insulin resistance and obesity.

Despite the presence of amyloid accumulation, tau pathology, and gliosis, DKI-lean mice performed similarly to WT controls in cognitive tasks. Meanwhile, DKI-HFHS mice showed spatial learning impairments despite possessing no greater AD-associated protein aggregates than the DKI-lean cohort. Body weight analysis confirmed significant differences between lean, standard, and HFHS-diet groups (Extended Data Fig. 11), underscoring that dietary composition can contribute to systemic metabolic stress and cognitive vulnerability without altering classical AD pathology. This reinforces the concept that brain insulin resistance contributes to AD-associated cognitive decline^14,53,54^. The results also highlight that dietary composition plays a crucial role in mitigating cognitive deficits associated with AD^28,55–57^, independent of the level of protein aggregate pathology. The DKI-HFHS model recapitulates a central feature of MetS—chronic inflammation—which, together with insulin resistance and vascular dysfunction, elevates AD risk^11,58^. Although the systemic nature of these symptoms is well characterized, the effects on brain physiology have been uncertain. Our data show that chronic dietary stress engages specific glial and neuronal programs that may underlie early cognitive decline, even in the absence of pronounced changes in amyloid or tau pathology. These findings support the hypothesis that MetS is not only a risk factor for AD but also a mechanistic contributor that reshapes cellular responses to neurodegeneration. The DKI-HFHS mouse model thus offers a valuable platform for investigating how systemic metabolic stress shapes glial and neuronal responses in the brain.

### Glial metabolic stress responses and microglia–synapse interactions

Recent studies have expanded beyond M1/M2 classifications to recognize a spectrum of microglial activation states associated with neurodegenerative diseases^33,59,60^. Using snRNA-seq, we identified a unique glial transcriptional program selectively induced in DKI-HFHS mice that we term the *<u>m</u>etabolic impairment in <u>n</u>euro<u>d</u>egeneration* (MinD) state. This state was most prominent in a microglial subcluster, MG3, defined by the selective upregulation of genes involved in synaptic regulation (e.g., Dlg2, Kcnip4, Lsamp, Ptprd, and Nrg3), transmembrane signaling, and amyloid pathology. Notably, the MinD signature is distinct from known DAM and inflammatory microglia profiles^33^ and absent in WT-HFHS mice, indicating that it represents a specific response arising from the intersection of diet-induced metabolic stress and AD pathology. The significant differences in body weight across diet groups (Fig. 1 and Extended Data Fig. 11) underscore how chronic dietary stress systemically alters metabolic physiology in ways that elicit distinct glial transcriptional states in the AD brain.

*In vivo*, we observed altered Trem2 expression and processing in DKI-HFHS microglia. Transcriptomic analysis revealed several MinD-upregulated genes, including Celf2 and Irf8, alongside downregulation of Inpp5d, all of which are known regulators of Trem2 signaling and processing. Celf2, an RNA-binding protein, regulates alternative splicing of Trem2 to produce soluble Trem2 (sTrem2), though its effects on sTrem2 abundance remain unclear^61,62^. Irf8, a transcription factor essential for microglial identity and activation, has been shown to regulate Trem2-dependent transcriptional programs and disease-associated microglia states^40,41^. Conversely, Inpp5d, a negative regulator of PI3K–AKT signaling downstream of Trem2, was downregulated, suggesting a shift toward enhanced Trem2 signaling responsiveness under metabolic stress^63^. Because sTrem2 can bind Aβ and influence its clearance or aggregation^38^, these transcriptomic changes may contribute to the altered Trem2 phenotype observed *in vivo*. This altered expression profile may, in turn, link diet-induced transcriptional shifts to alterations in Aβ dynamics under metabolic stress.

Among microglial MinD genes, Nrg3 emerged as a candidate linking glial transcriptional shifts to inhibitory synaptic signaling. Nrg3 is a selective ligand for ErbB4, an inhibitory synapse-enriched receptor^45,64,65^. ErbB4 signaling itself has been linked to metabolic dysregulation and clinical obesity^66^. In the DKI-HFHS cortex, Nrg3 protein levels were increased, suggesting enhanced microglia-derived Nrg3-ErbB4 engagement. Indeed, inhibitory synapse density was substantially reduced. We also observed Nrg3 upregulation in astrocytes and oligodendrocytes, indicating that multiple glial populations contribute to ErbB4-mediated signaling. We hypothesize that the MinD transcriptome in glia alters inhibitory circuit integrity at least in part through Nrg3-mediated mechanisms.

While our *in vivo* validation of MinD focused on microglia, the astrocytes and oligodendrocytes of DKI-HFHS mice also exhibited MinD-like transcriptional changes. Previously, we reported that the transcriptional responses of astrocytes and oligodendrocytes follow delayed inflammatory trajectories compared to microglia in the DKI strain, becoming more pronounced at 20 months than at 10 months^26^. The emergence of a robust glial-wide MinD state at 10 months suggests an earlier and coordinated adaptation to metabolic stress across glial populations. Therefore, a glia-wide MinD state may represent an early step in the pathological cascade linking MetS-driven metabolic dysfunction to AD progression. Surprisingly, several glial MinD state genes (e.g., Lrrtm4, Dlg2, and Ptprd) are risk factors for autism spectrum disorders^67,68^, another condition associated with metabolic dysfunction^69^. This highlights the potential broader significance of the glial MinD states in other metabolic disorders.

### Meis2 upregulation in inhibitory neurons under metabolic stress

One of the most striking findings from our study was the selective upregulation of the transcription factor Meis2 in L2/3 inhibitory neurons in HFHS-fed mice, with further increased expression in DKI-HFHS mice. This robust diet-specific response was absent in DKI-Lean mice, an indication that Meis2 acts as a metabolic stress– responsive factor that is further dysregulated in the AD context. Meis2 is involved in glucose homeostasis in pancreatic β-cells, forming transcriptional complexes with Pbx1 and Pbx3^70–73^. Remarkably, these same genes are among the top five cell identity markers in Meis2^+^ L2 inhibitory neurons. These findings suggest that Meis2^+^ neurons may engage a metabolic homeostasis program in response to dietary stress, paralleling its transcriptional role in pancreatic β-cells as a mitigator of hyperglycemic signals. Other top genes enriched in this cell population include Auts2, an autism risk gene^74^, and Kcnb2, which regulates both insulin secretion^75^ and neuronal excitability^76,77^.

Meis2^+^ inhibitory neurons exhibited transcriptomic enrichment for pathways regulating vesicle release, synaptic organization, and membrane excitability, which are selectively altered under HFHS-diet conditions. In contrast, adjacent L2-4 excitatory neurons exhibit distinct transcriptional changes in response to disease and diet, but converge on similar signaling cascades, all of which are linked to synaptic signaling. Whether these changes are compensatory or pathogenic remains unclear, but they indicate focal changes in L2-3 cortical physiology driven by altered inhibition and merit further investigation.

### Transcriptomic consequences of insulin signaling and resistance

Metabolic syndrome, with or without AD, has been associated with impaired brain insulin signaling, including disruptions in IRS-1 and AKT pathways, indicative of a brain insulin-resistant state^13–15,54,78^. These disruptions have motivated clinical trials of central insulin delivery as a therapeutic strategy for AD^16,79^. In contrast, our study focused on the transcriptional consequences of insulin resistance. We observed pronounced gene expression changes across glial types and within a specific class of inhibitory neurons in response to metabolic stress. Future work will determine whether these transcriptional shifts are driven by canonical insulin signaling pathways or mediated through alternative mechanisms linking metabolic dysfunction to gene regulatory networks.

## Conclusion

By comparing models of impaired insulin production and systemic resistance, we identified how these distinct metabolic disruptions contribute to neuroinflammation, synaptic dysfunction, and cognitive decline. Our findings demonstrate that chronic dietary stress induces cognitive impairments and transcriptional alterations in AD-vulnerable brain regions via mechanisms not solely attributable to hyperglycemia or classical AD pathology. The emergence of a shared glial MinD program across microglia, astrocytes, and oligodendrocytes, along with parallel transcriptional changes in Meis2^+^ inhibitory neurons, underscores the multifaceted impact of metabolic stress on cortical circuitry. The absence of overt changes in AD pathology in the HFHS-fed mice is notable given the apparent cognitive deficits and transcriptional alterations. These results suggest that MetS reshapes the cellular landscape of the AD brain in ways not captured by conventional neuropathology. A limitation of this study is the absence of direct assessment of L2-4 circuit function as a result of the glial and neuronal expression changes. We hypothesize that these collective cellular alterations result in cognitive impairment in DKI-HFHS mice. Clinically, MetS has been associated with worsened outcomes in early-stage AD and paradoxical protection later in disease progression^80,81^, suggesting a dynamic interplay between metabolic and neurodegenerative processes. Future studies will investigate whether glial MinD states are neuroprotective, maladaptive, or a mixture, and whether their function shifts during disease progression. By modeling the compound effects of diet and disease, this study provides a tractable framework for understanding how metabolic dysfunction influences age-related neurodegeneration and identifies novel cellular targets for therapeutic intervention.

## Methods

### Animal handling and husbandry

Studies were approved by Yale University”s Institutional Animal Care and Use Committee (IACUC, protocol #07281). Mice were cared for by the Yale Animal Resources Center (YARC) and housed in groups of one to five mice per cage with a 12-hour light/dark cycle, with food and water available *ad libitum. App*^*NL-F*^*/Mapt*^*hMAPT*^ and age-matched wild-type C57BL/6J (WT) mice were used for the streptozotocin (STZ) treatment study. *App*^*NL-F/NL-F*^ mice were generated as previously described^22,23^, then crossbred with *Mapt*^*hMAPT/hMAPT*^ mice^24^. Double knock-in *App*^*NL-G-F*^*/Mapt*^*hMAPT*^ (DKI) and age-matched WT mice were used for the diet-induced glucose intolerance resistance studies. DKI mice were generated by crossbreeding *Mapt*^*hMAPT/hMAPT*^ and *App*^*NL-G-F*^ single knock-in mice as previously described^25,26^. Sex was balanced across all cohorts, with equal numbers of male and female mice per group. Single knock-in *App*^*NL−F/NL−F*^, *App*^*NL−G−F/NL−G−F*^, and *Mapt*^hMAPT/hMAPT^ mice were generously provided by Drs. Saito and Saido, RIKEN Center for Brain Science. All mouse lines used were maintained on a C57BL/6J background after more than nine generations of backcrossing.

### STZ-Induced Type 1 Diabetes Model

To model Type 1 diabetes, we utilized a multiple low-dose streptozotocin (STZ; Sigma, S0130) administration protocol^19,82,83^. *App*^*NL-F*^*/Mapt*^*hMAPT*^ and WT mice were aged to 8 months. STZ was freshly prepared in 50 mM sodium citrate buffer (pH 4.5) at a final concentration of 6 mg/mL and used within 10 minutes of preparation. On the first day of STZ treatment, all food was removed from cages 4 hours prior to injection, while mice continued to have access to regular drinking water. Mice were weighed and randomized into experimental groups with balanced genotype and sex distributions (2 cohorts, *n=15-20* mice/group). STZ was administered via intraperitoneal (i.p.) injection at a dose of 40 mg/kg. Vehicle (Veh) control mice received an equal volume of sodium citrate buffer. Following injection, mice were returned to their cages and provided with standard chow and 10% sucrose water. STZ injections were administered once daily for five consecutive days at a dose of 40 mg/kg. On the final day of STZ treatment, the 10% sucrose water was replaced with regular drinking water. Mice had *ad libitum* access to food and water for the remainder of the study. Mice with blood glucose levels below 200 mg/dL, 7 weeks after the initial STZ series, received a second 5-day course of 40 mg/kg i.p. injections, following the same protocol, with standard food and regular drinking water provided.

### HFHS-Induced Type 2 Diabetes Model

To model Type 2 Diabetes, a chronic high-fat-high-sugar (HFHS) diet was used ^20^. DKI and WT mice were randomized into aged and sex-balanced groups (*n=15-20* mice per group), initially fed *ad libitum* on a standard chow diet (Tekland Global 18% protein diet, S2018). At approximately 3-4 months of age, mice were switched to a lean diet (Research Diets, D12450K; diet composition: protein 20%, fat 10%, and sucrose 0% kcal) or HFHS diet (Research Diets, D12492; diet composition: protein 20%, fat 60%, and sucrose 275% kcal). Mice were maintained on each respective diet until the date of sacrifice.

### Assessment of metabolic alterations

Body weight and fasting blood glucose levels were used to assess the metabolic state of mice in each diabetic model. STZ-treated mice were weighed at 0, 2, 7, and 18 weeks post-injection (Extended Data Fig. 1a). Mice on lean or HFHS diets were weighed every three weeks, starting one month after diet onset (Fig. 1a-c). For blood glucose measurements, mice were fasted for 6 hours, and blood glucose levels were subsequently measured from the tail vein using the OneTouch Basic (LifeScan, Inc., USA) blood glucose monitoring system. Mice were considered hyperglycemic if their blood glucose levels exceeded 150 mg/dL, and classified as diabetic if levels exceeded 300 mg/dL, in accordance with established guidelines (Furman, 2021). An oral glucose tolerance test (GTT) was used to assess the acute metabolic response to glucose (Andrikopoulos et al., 2008; Ayala et al., 2010) (Fig. 1e, Extended Data Fig. 1c). Mice were fasted for 6 hours and then administered a bolus of 20% dextrose solution (2 g/kg body weight) via oral gavage. Blood glucose levels measured 1 minute before and 30 minutes after dextrose administration were used to compare metabolic impairment between groups.

### Morris Water Maze

Spatial learning and memory were assessed using the Morris Water Maze (MWM), conducted as previously described with slight modifications ^23,25,26^. Briefly, a 1m diameter pool, surrounded by visual cues, was filled with water and maintained at a temperature between 21-23°C. An invisible platform was submerged 1.5 cm underwater in the target quadrant. All visual cues, water temperature, and ambient lighting were kept constant throughout the experiment. Mice underwent 4 days of habituation training and were then randomized for MWM assessments with genotypes blinded to the experimenter. Testing occurred over eight days: three days of forward training, 1 probe day for the forward acquisition paradigm, immediately followed by 3 days of reverse training and 1 probe day for the memory retrieval paradigm. All mice within a single cohort were tested at the same time. Each acquisition training session included four swim trials conducted twice daily, once in the morning and again in the afternoon, totaling eight trials. Mice were randomly placed in one of four entry points, facing away from the pool”s center and opposite the target quadrant. Entry locations were varied for each mouse in each training session. Each trial lasted for a maximum of 60 seconds, with success defined as the mouse locating and remaining on the platform for at least 1 second. In the initial training session, if a mouse failed to find the platform within the time limit, it was guided backwards to the invisible platform and allowed to rest there for 10 seconds before being removed from the pool. The probe trial took place 24 hours after the final training session, during which the transparent platform was removed, and mice were placed facing the wall opposite the target quadrant and allowed to swim for 60 seconds. The reverse training sessions and probe trial followed the same protocol, but the transparent platform was placed in the opposite quadrant. Mouse latencies to the platform were recorded using a JVC Everio G-series camcorder (JVCKenwood, Yokohama, Japan) and tracked by Panlab”s Smart software (Harvard Bioscience, Massachusetts, USA). Following the final probe trial, mice were assessed for visual acuity deficits. A visible platform was positioned in the target quadrant, and trials continued until latencies stabilized over a maximum of 15 trials, with average latencies calculated for the final three trials. Mice in the STZ-treatment study underwent MWM evaluations at 12 months (Extended Data Fig. 1), 17 weeks post-STZ-treatment, while mice in the HFHS diet study were evaluated at 10 months (Fig. 1 and Extended Data Fig. 2).

### Immunolabeling of mouse tissue

Mice were euthanized with CO_2_ and perfused transcardially with ice-cold phosphate-buffered saline (PBS, pH 7.4), followed by ice-cold 4% paraformaldehyde (PFA) in PBS. Brains were rapidly dissected, bisected at the midline, and post-fixed in 4% PFA at 4°C for 24 hours. Post-fixed brains were sectioned into 50 μm coronal slices using a Leica VT1000S vibratome (Leica, Wetzlar, Germany). Free-floating sections were stored long-term at 4 °C in PBS containing 0.05% sodium azide. For immunofluorescence labeling, free-floating sections were incubated at room temperature for 1 hour in blocking buffer (5% bovine serum albumin, 1% Triton-X, pH 7.4 PBS), then incubated overnight at 4°C in primary antibodies diluted in blocking buffer.

The following primary antibodies were used: rabbit anti-ß amyloid (1:600, RRID:AB_2797642) for Ab deposition; rabbit anti-IBA1 (1:600, RRID:AB_839504), goat anti-IBA1 (1:500, RRID:AB_10982846), rat anti-Cd68 (1:250, RRID:AB_322219) sheep anti-Trem2 (1:125, RRID:AB_356109), and goat anti-Nrg3 (1:300, RRID:AB_2154802) for assessment of microgliosis; chicken anti-GFAP (1:1000, RRID:AB_304558) for astrocyte gliosis; rabbit anti-p(Y128)ErbB4 (1:200, RRID:AB_940986), guinea pig anti-Gephyrin (1:200, RRID:AB_2661777), guinea pig anti-Homer (1:200, RRID:AB_2661777), rabbit anti-PSD-95 (1:300, RRID:AB_87705), chicken anti-Synapsin½ (1:250, RRID:AB_2622240), rabbit anti-VGAT (1:200, RRID:AB_887871), and rabbit anti-VGLUT1 (1:200, RRID:AB_887877) for synaptic density measurements; rabbit anti-Meis2 (1:250, RRID:AB_11026529) or rabbit anti-PBX1 (1:250, RRID:AB_11205760) co-labeled with chicken anti-NeuN (1:400, RRID:AB_11205760) for the assessment of Layer 2 (L2) inhibitory neurons; guinea pig anti-Pvalb (1:700, RRID:AB_2156476), WFA-FITC (1:2000, Vector Laboratories, FL-1351-2), mouse biotinylated AT8 (1:100, RRID:AB_223648), and rabbit anti-pThr217 (1:200, RRID:AB_2533741) for assessment of Tau pathology. For optimal immunodetection of AT8, the Alexa Fluor™ 488 Tyramide SuperBoost Kit, Streptavidin (Invitrogen, B40932), was used following the manufacturer”s protocols. For pThr217 staining, an antigen retrieval step was performed prior to exposure to primary antibody by incubating slices in 1x Reveal Decloaker buffer (RV 1000 M, Biocare Medical) for 15 min at 90°C in an incubator and then at room temperature for 15 minutes.

Sections were washed 3 times for 5 min each in blocking buffer and incubated in Alexa Fluor^TM^ secondary antibodies diluted in blocking buffer (donkey or goat anti-chicken, anti-goat, anti-guinea pig, anti-rabbit, anti-rat, 1:200, ThermoFisher Scientific) for one hour at room temperature in the dark. To reduce background autofluorescence, sections were washed three times in PBS, briefly dipped in ddH_2_O, and incubated for one minute in CuSO_4_ (10 mM CuSO_4_ in 50 mM ammonium acetate buffer, pH 5.0) followed by a second brief dip in ddH_2_O. Sections received a final series of three PBS washes for 5 minutes, then were mounted onto Superfrost Plus microscope slides (ThermoFisher Scientific, 22-037-246) and coverslipped with Prolong^TM^ Glass mounting medium (ThermoFisher Scientific, P36984).

### Imaging and analysis of mouse tissues

All images were acquired using the Stellaris STED 8 confocal microscope (Leica Microsystems, Wetzlar, Germany). Tissues immunolabeled to assess for Aβ deposition, microgliosis (Cd68, Iba1, Trem2, Fig. 2), astrogliosis (GFAP), Tau pathology (AT8, pThr217), perineuronal nets (Pvalb and WFA), and the expression of Meis2 in NeuN^+^ cells, were imaged with a 20X, 0.8NA air objective (Fig. 2, and Extended Data Fig. 7 and 9). Images were acquired as z-stack images with a 2 μm step size within the motor and somatosensory cortices, specifically targeting L2-4. All images were analyzed as maximum intensity projections, and the percent area occupied by threshold immunolabeled signal was quantified using ImageJ (v2.16.0, License GPLv3+). Statistical analysis was performed at the level of individual mice using the average of two sections per mouse. Binary images were generated from NeuN and Meis2 fluorescence channels. For Meis2^+^ cell quantification, A NeuN-based mask was used to identify neuronal nuclei, and Meis2+ signals within this mask were isolated using *the Image Calculator* function in ImageJ. The number of Meis2+ cells was quantified using *Analyze Particles* and expressed as a percentage of total NeuN+ cells (Fig. 8). Quantification was performed within a 200 × 500 μm region of cortical layer 2, using single z-stack planes averaged across all planes in the stack. Two sections were analyzed per mouse, and statistical comparisons were performed at the animal level.

Microglial morphology and Aβ plaque association were assessed from IBA1 and Aβ co-immunolabeled images acquired within cortical layers 2-3. Full-depth z-stacks, with 1 μm step size, were collected using a 40X, 1.3 NA oil-immersion objective with 2X digital zoom, with 5 to 6 regions imaged across two sections per mouse. To quantify Aβ plaque volume and microglia–plaque associations, 3D volumes were generated from the IBA1 and Aβ channels using the *Surfaces* tool. Aβ plaques were defined as objects larger than 5 μm^3^. Using the object-to-object statistics feature, the number of IBA1^+^ microglial surfaces in direct contact with Aβ plaques was quantified and reported as a proportion of total microglia per image (Extended Data Fig. 4). For microglial Sholl analysis, the *Mask* tool was used to select and isolate the IBA1^+^ signal within the volume. Then, the *filament* tool was used to trace microglial processes within the masked region for morphological reconstruction and Sholl quantification (Extended Data Fig. 3). Between 4 to 6 microglial cells were analyzed per mouse.

For synaptic density quantifications, synaptic proteins were labeled with STED-compatible secondary antibodies: Alexa Fluor^TM^ 488 and Alexa Fluor ^TM^ 555 (1:200, ThermoFisher Scientific). Z-stack images were acquired with a 100X, 1.4 NA oil-immersion objective with 2X digital zoom and a 0.5 μm step size. Dual-channel imaging was performed with a pulsed White Light Laser for excitation and 660-nm depletion laser beams. Synaptic loci were quantified in Imaris using the *Spots* tool to detect pre- and post-synaptic signals. Colocalization was defined as pre- and post-synaptic spots located within 0.4 μm of each other in three-dimensional space.

### Single-nuclei RNA sequencing and analysis

HFHS cohort mice were sacrificed at 11.5 months by rapid decapitation. Brains were harvested and medially dissected on ice using a scalpel. Brain hemispheres were micro-dissected to isolate the cortex and hippocampus, then immediately frozen on dry ice and stored at −80 °C. For single-nucleus RNA sequencing experiments, cortical and hippocampal tissues from the left-brain hemisphere of a single mouse were pooled together. Preparation of samples, cDNA library construction, and sequencing were performed as previously described ^26,31^. Briefly, pooled cortical and hippocampal tissues were gently homogenized on ice in a buffered solution (10 mM HEPES, pH 7.9; 25 mM KCl; 1 mM EDTA; 10% glycerol; 2 M sucrose) supplemented with 80 U/mL RNase inhibitor (Roche 03335402001) to release cellular nuclei. Samples were centrifuged at 20,000 x g at 4 °C for 1 hour, supernatants were discarded, and pelleted nuclei were resuspended to a concentration of 700-1200 nuclei/mL in chilled PBS supplemented with 80 U/mL RNase Inhibitor. Nuclei barcoded cDNA libraries were constructed using the Chromium Single Cell 3” Reagents Kit v3 (10x Genomics) following the manufacturer”s guidelines. Sample libraries were pooled and collectively run on an Illumina NovaSeq 5000 using single-indexed paired-end HiSeq sequencing, achieving a mean sample sequencing depth of 450 million reads and 30,000 UMIs per nucleus (Extended Data Figure 5). Raw sequencing reads were mapped against the mm10-2020-A mouse reference gene using the Cell Ranger Count pipeline (v6.1.2, 10x Genomics) with default parameters.

Processing of snRNA-seq data was conducted as previously described, with modifications^26^. Sample gene count matrices were merged into a single AnnData object and processed using the Scanpy v1.9.6 Python package^84^. Quality control filtering removed nuclei with more than 5% of transcripts mapping to mitochondrial genes, fewer than 50 detected genes, or more than 5,000 genes. Genes expressed in fewer than 10 nuclei were also excluded. Clustering was performed in two stages. First, unsupervised clustering was conducted using Scanpy for the unbiased identification and removal of putative doublets. Gene expression data were normalized and log1p-transformed. Highly variable genes were identified using *scanpy*.*pp*.*highly_variable_genes* with *n_top_genes=2000* and *flavor=“seurat_v3”*. Principal component analysis of variable genes was computed with *scanpy*.*tl*.*pca*, specified with *svd_solver=“arpack”*. Using the top 37 principal components, a neighborhood graph was constructed using scanpy.pp.neighbors, followed by functions *scanpy*.*tl*.*leiden* for Leiden Clustering^85^ and *scanpy*.*tl*.*umap* for UMAP embedding. Cluster identities were initially assigned based on marker genes identified in our previous studies^26,31^. Doublets, defined as individual clusters enriched for marker genes from multiple cell types, were filtered from the dataset.

Following doublet removal, downstream analyses were performed using the *model*.*SCVI* (scvi-tools v0.20.3)^86^. The model was trained on the filtered dataset using total gene counts and percent mitochondrial content as covariates. The latent representation from the trained scVI model was then used for batch-corrected dimensionality reduction, clustering, and visualization in UMAP space. Cluster classification and subclass annotation were based on the union of known marker gene expression profiles and alignment to the Allen Brain Cell Atlas using MapMyCells (RRID:SCR_024672) (Supplementary Table 2).

To achieve higher-resolution analysis, specific cell populations (microglia, excitatory neurons, inhibitory neurons) were subset from the full dataset and re-analyzed independently. For each subset, a new scVI model was trained *de novo* on the cell-type-restricted AnnData object to account for population-specific variation. The latent space inferred from the scVI model was then used to perform batch-corrected reclustering and visualization (Figs. 5, 7, and 9, and Extended Data Fig. 5–7).

### Differential expression and pathway enrichment analysis

Differential expression gene (DEG) analysis was performed using the *scanpy*.*tl*.*rank_genes_groups* function with the Wilcoxon rank-sum test, comparing experimental groups within each identified cell type. DEGs were defined as those expressed in at least 10% of nuclei tested, with an adjusted *P*-value (Benjamini-Hochberg correction) less than 0.05, and an absolute log-transformed fold change greater than 0.25. Gene pathway enrichment and cellular compartment analysis were conducted using Cytoscape (v3.9.1)^87^ with the ClueGO (v2.5.9)^88^ and STRING (v2.0.0)^89^ plugins. Ingenuity Pathway Analysis (IPA, Qiagen) was used to determine the subcellular localization of DEGs.

### Statistics and reproducibility

No statistical methods were used to predetermine sample sizes, but all experiments were conducted with comparable sample sizes. Data collection and downstream analyses were performed by experimentalists blinded to experimental conditions, and automated where possible to minimize experimenter bias. Data were analyzed using GraphPad Prism v9.2.0. Experimental analysis involving more than two groups was evaluated using two-way ANOVA, followed by appropriate multiple comparison tests. Normality tests were performed to determine the correct application of parametric or non-parametric methods. Unless indicated otherwise, data are presented as mean ± SEM, with individual data points representing intra-mouse means. Statistical details, including the exact *number (n) of* biological replicates (animals or cells), numerical means, statistical tests, and multiple comparison corrections, are provided in the figure legends. Statistical significance is denoted as \**p<*0.0332, \*\**p<*0.0021, \*\*\**p<*0.0002, and \*\*\*\**p<*0.0001.

## Data availability

Raw FASTQ sequencing files and gene count matrices have been deposited in NCBI”s Gene Expression Omnibus (GEO) under accession code GSE262426.

## Acknowledgements

We acknowledge the Yale Pathology Department and the Center for Human Brain Discovery, which is supported by philanthropic funds, for providing human brain and non-CNS tissues.

## Funding

This work was supported by an award from the Kavli Institute for Neuroscience at Yale University to L.N. and by grants R01AG034924, P30AG066508, R01AG070926 from the NIA to S.M.S.

## Author contributions

Conceptualization, L.N. and S.M.S.; Methodology, L.N., J.T., T.K., H.A. and S.M.S.; Investigation, L.N., J.T., T.K., H.A.; Writing – Original Draft, L.N. and S.M.S.; Writing – Review & Editing, all; Funding Acquisition, L.N. and S.M.S.; Resources, S.M.S; Supervision, S.M.S.

## Ethics declarations

The author declare no competing interests.

## Extended data figures

**Extended Data Fig. 1.**
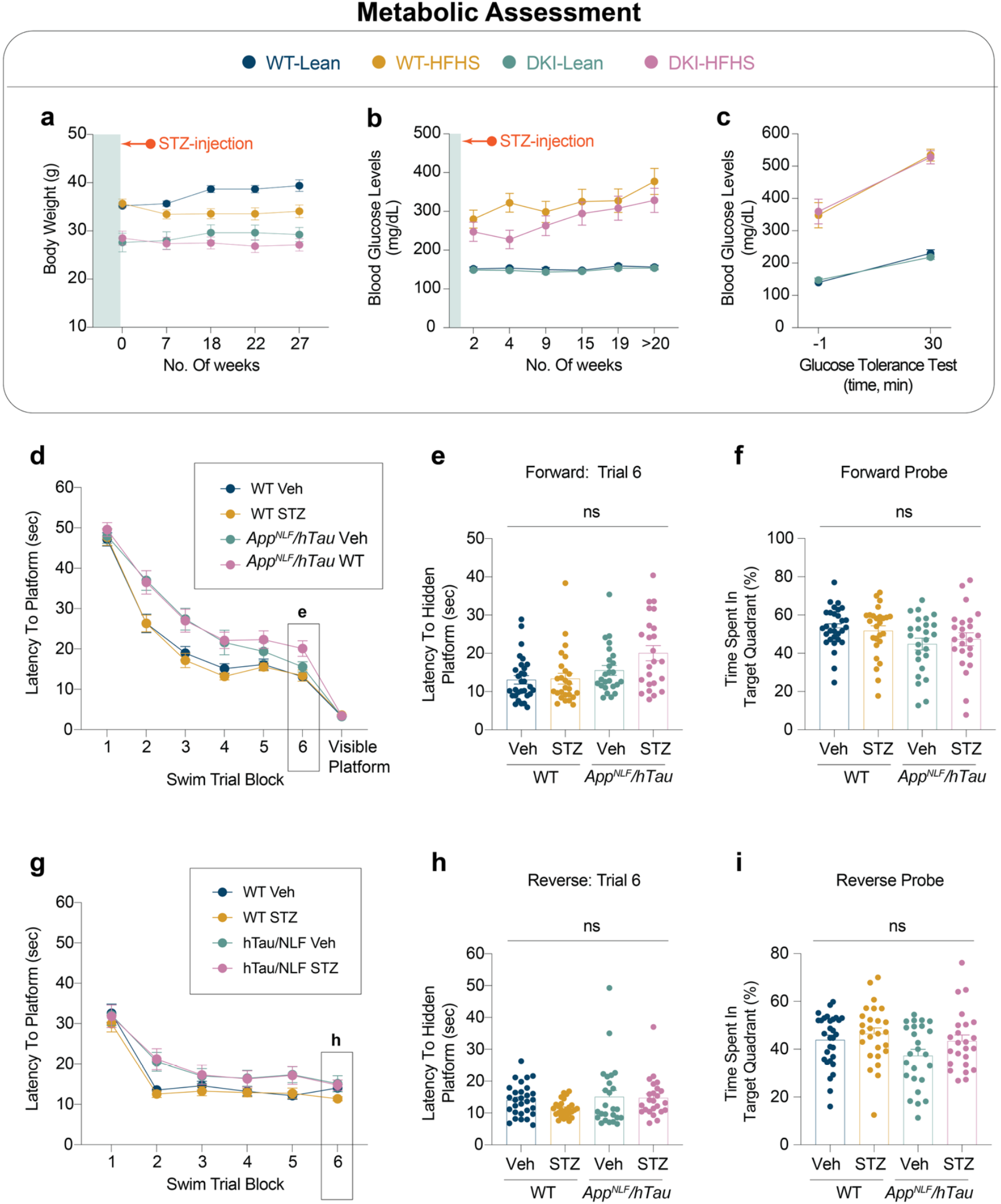
Insulin deficiency induced by STZ does not lead to obesity or cognitive deficits. **a-c**, Metabolic assessments following STZ treatment. Body weight (**a**), Blood glucose level measurements (**b**), and glucose intolerance test 30 minutes post oral glucose bolus delivery (**c**). **d-i**, MWM performance evaluation across acquisition (forward training, **d-f**) and reversal (reverse training, **g-i**) sessions. **d-e**, Latency to the hidden platform during the training phase (**d**) and final training session (**e**). **f**, Fraction of time spent in the target quadrant during acquisition probe trials. **g-h**, Latency to the hidden platform during the reverse training phase (**g**) and final reverse trial (**h**). **i**, Fraction of time spent in the target quadrant during the reverse probe trial. (**a-g**, *n=31* for WT-Veh, *n=25* for WT-STZ, *n=25* for *App*^*NLF/hTau*^*-*Veh, and *n=24* for *App*^*NLF/hTau*^*-*WT). Data are presented as mean ± SEM and analyzed by two-way ANOVA with Tukey”s multiple comparisons test. STZ, streptozotocin; ns, not significant.

**Extended Data Fig. 2.**
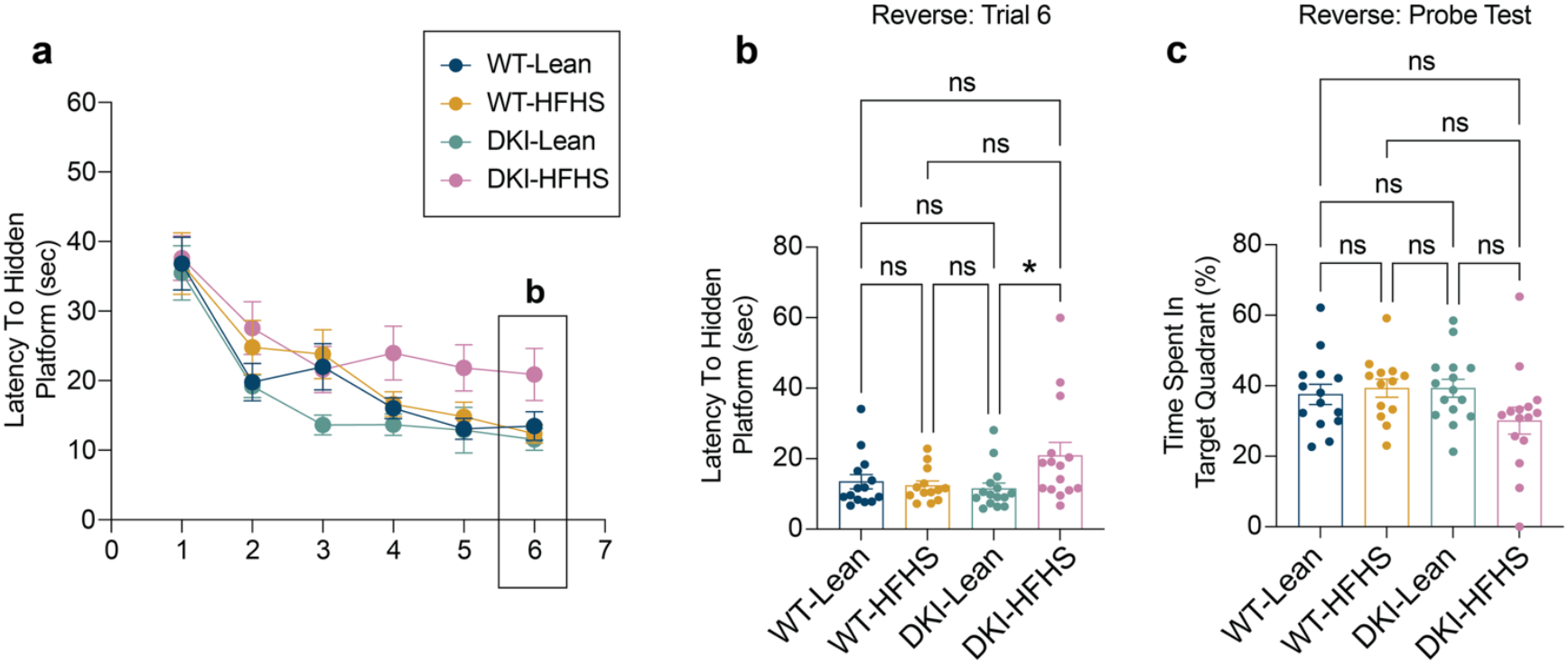
DKI-HFHS mice experience deficits in learning, but not memory consolidation. **a-c**, Reverse learning performance in the MWM. **a-b**, Latency time to the hidden platform during the reverse training phase (**a**) and final training session (**b**). **c**, Fraction of time spent in the target quadrant, during reverse probe trials. (*n=14* for WT-Lean and WT-HFHS groups; *n=15* for DKI-Lean and DKI-HFHS groups). Data are presented as mean ± SEM and analyzed by two-way ANOVA with Tukey”s multiple comparisons test, \**p<*0.0332, \*\**p<*0.0021, \*\*\**p<*0.0002, and \*\*\*\**p<*0.0001; *ns, not significant*. Data related to Fig. 1.

**Extended Data Fig. 3.**
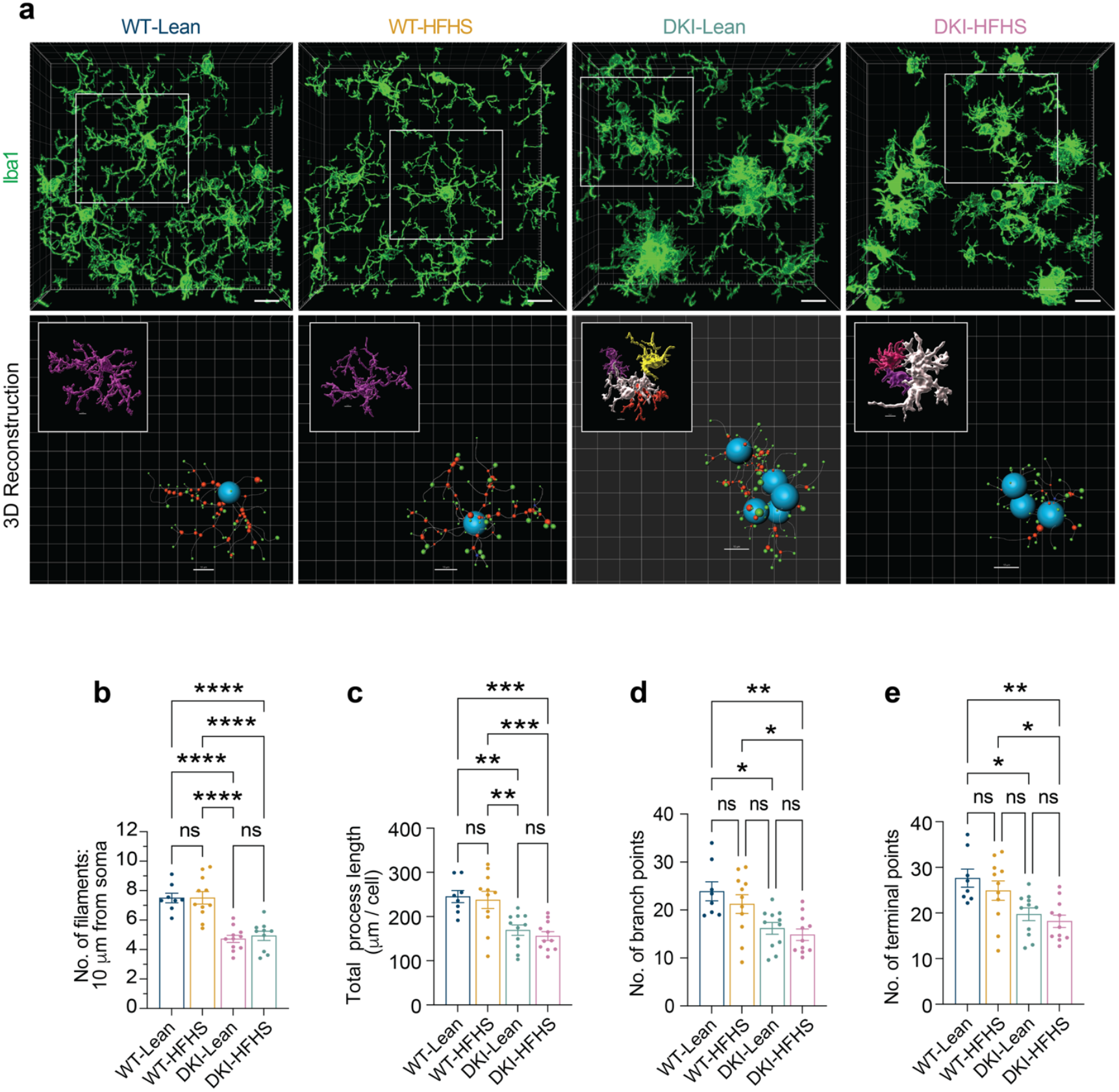
Filament tracing and Sholl analysis of microglial complexity. **a-e**, Assessment of microglial morphology using filament tracing in Imaris. Data related to Fig. 2. **a**, Representative 3D renderings of microglia morphologies generated in Imaris (top) and corresponding filament tracings of the boxed region (bottom). Soma are shown in blue, branches in red, and terminal points in green. **b**, Quantification of the number of microglia filaments at a radial distance of 10 μm from the soma. **c-e**, Quantification of total process length (**c**), number of terminal points (**d**), and number of branch points (**e**). Data are presented as mean ± SEM and analyzed by two-way ANOVA with Tukey”s multiple comparisons test, \**p<*0.0332, \*\**p<*0.0021, \*\*\**p<*0.0002, and \*\*\*\**p<*0.0001; *ns, not significant*. Data related to Fig. 2.

**Extended Data Fig. 4.**
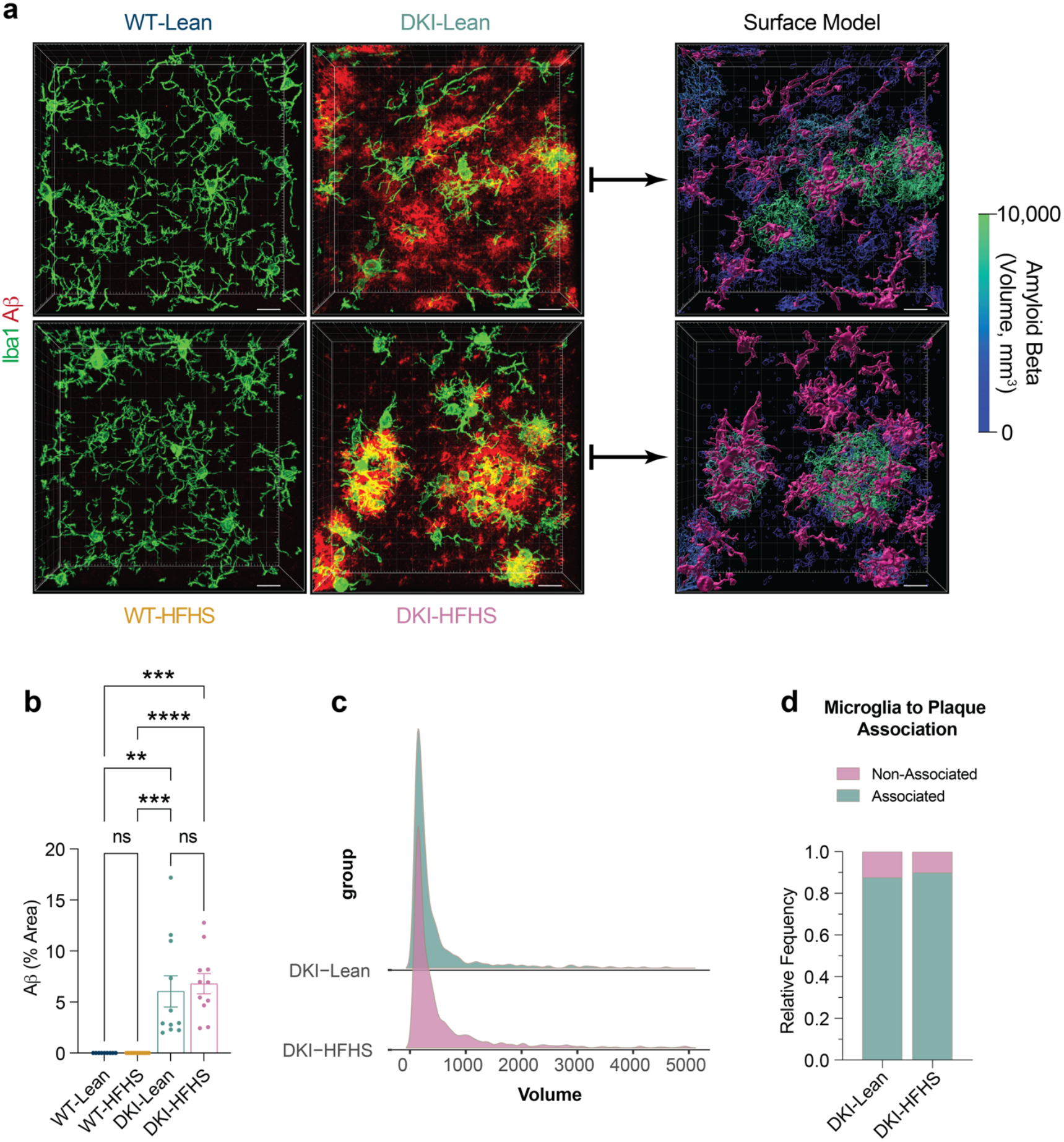
Aβ pathology and microglial plaque association are not impacted by HFHS diet. **a-d**, Assessment of microglial morphology and plaque associations using volumetric reconstructions in Imaris. **a**, Representative 3D renderings of microglial morphology (Iba1, green) and amyloid beta deposition (red). Example Imaris-generated surface models of microglia and amyloid plaques for DKI groups used in subsequent analyses. **b**, quantification Aβ deposition shown in **a. c**, Distribution of amyloid beta plaque volumes. **d**, Frequency of microglia-plaque associations. Data related to Fig. 2 and Extended Data Fig. 5a-d.

**Extended Data Fig. 5.**
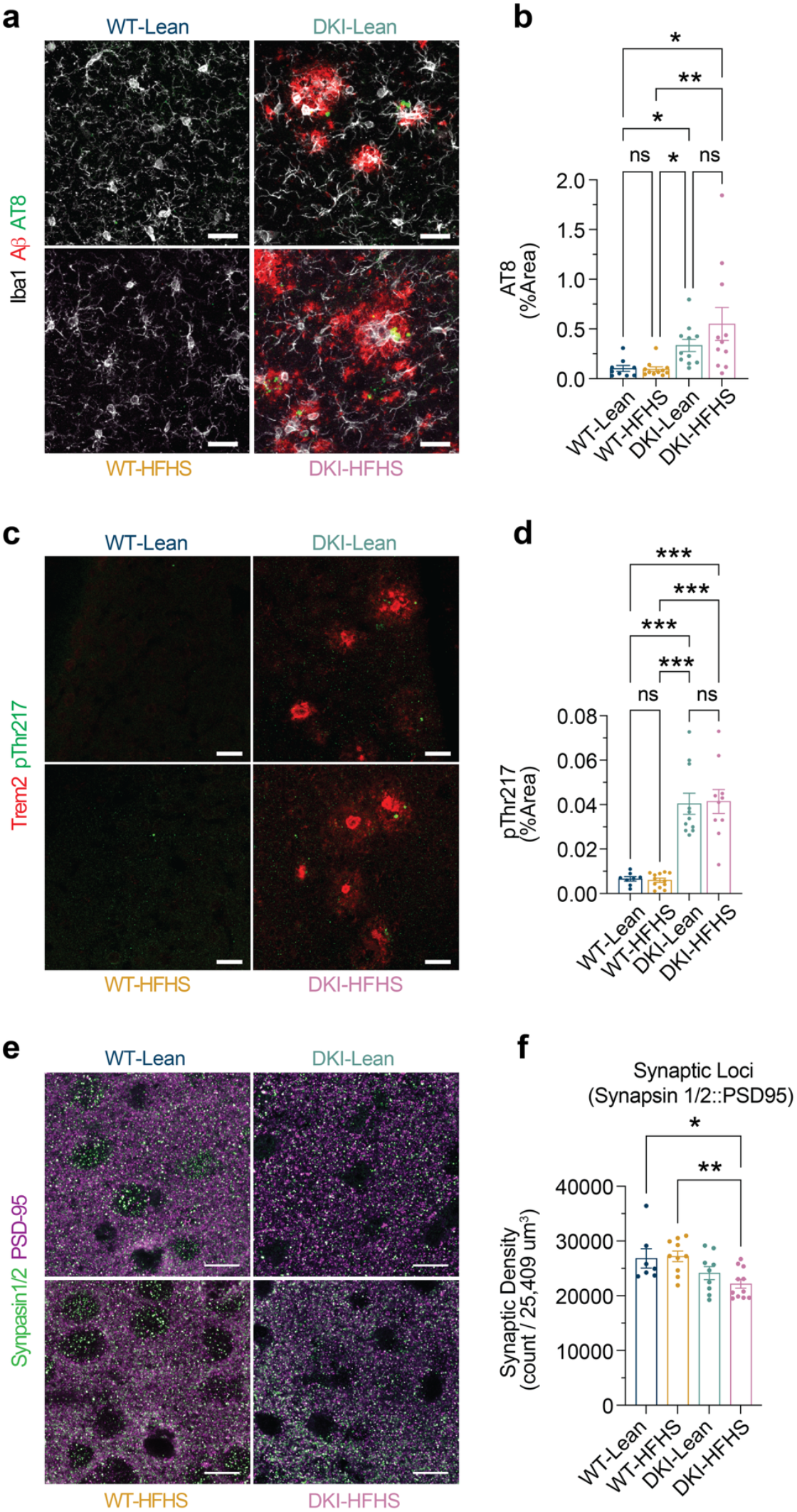
HFHS-diet induced synaptic deficits occur independently of tau pathology in DKI-mice. **a-d**, Image analysis of Tau makers AT8 and pThr217. **a**. Representative images of anti-Iba1 (white) and anti-Aβ (red), anti-AT8 (green) immunostaining across treatment groups. **b**, Quantification of AT8 percent area represented in **a**. Each data point represents the mean value of three images per mouse. **c**, Representative images of anti-Trem2 (red), anti- pThr217 (green) immunostaining. **d**, Quantification of pThr217 percent area represented in **c**. Each data point represents the mean value of 4-6 images per mouse. **e-f**, Representative staining of pre-synaptic anti-Synapsin1/2 and post-synaptic PSD-95 colocalization within cortical region L2 (**e**), synaptic quantification (**f**). Scale bars, 10 μm. Data related to Fig. 4. (**b**,**d**,**f**, *n=9* for WT-Lean group; *n=12* for WT-HFHS, *n=11* for DKI-Lean, and *n=10* for DKI-HFHS groups). All data presented as the mean ± SEM and analyzed by two-way ANOVA with Kruskal-Wallis test with Dunn”s multiple comparisons corrections (**b**,**d**) or Tukey”s multiple comparisons test (**f**). \**p<*0.0332, \*\**p<*0.0021, \*\*\**p<*0.0002, and \*\*\*\**p<*0.0001; ns: not significant. (*n=6*, WT-Lean; *n=10*, WT-HFHS; *n=10*, DKI-Lean; and *n=11*, DKI-HFHS groups).

**Extended Data Fig. 6.**
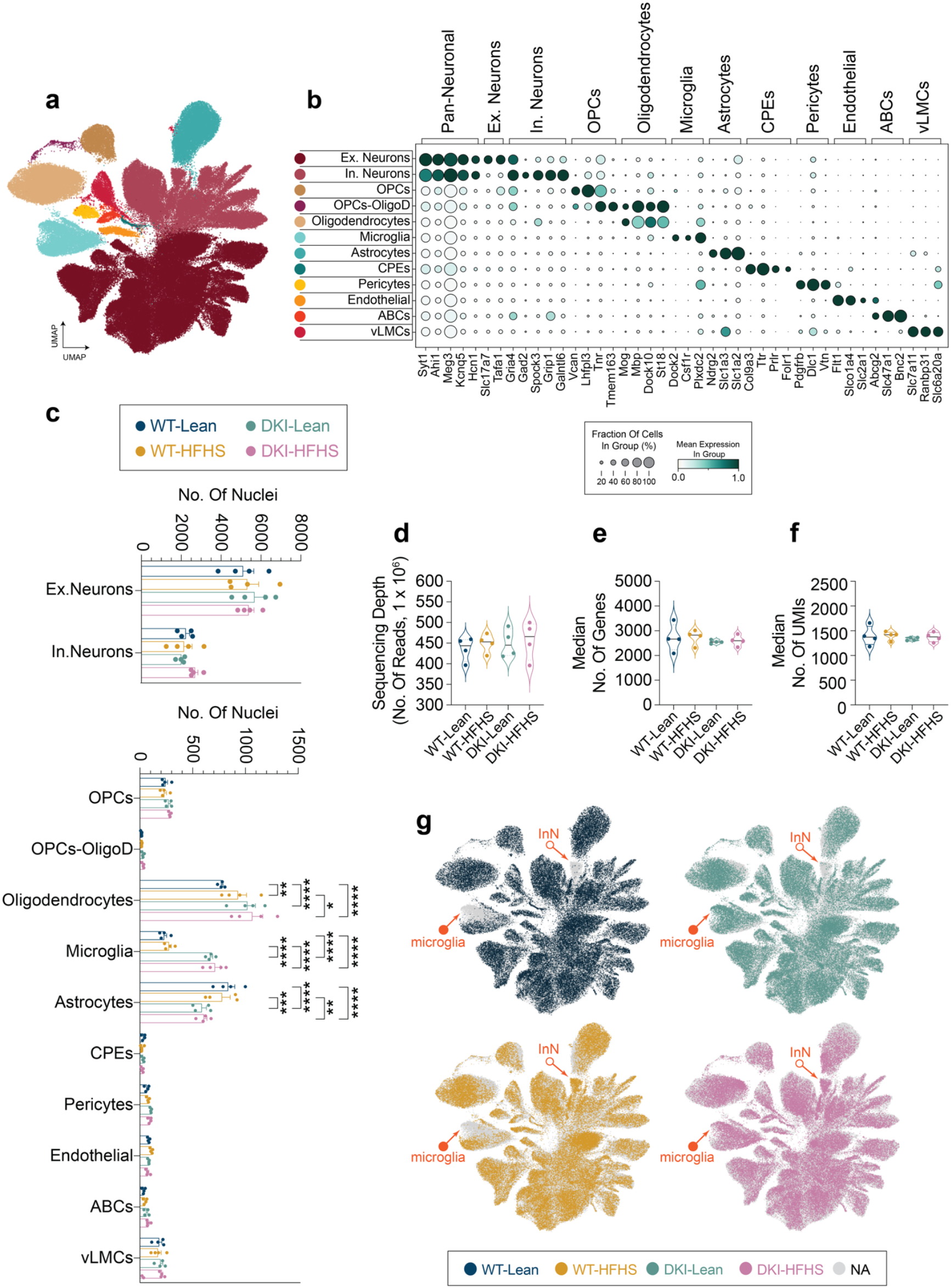
Single-nucleus RNA-seq reveals prominent transcriptional shifts in glial and inhibitory neuron populations. **a-g**, Integration and cell type classification of snRNA-seq datasets (n = 4 per group). **a**, UMAP embedding of single-nucleus transcriptomes showing cellular clustering based on gene expression profiles. Each point represents a single nucleus colored by annotated cell types shown in **b. b**, Dot plot showing the percentage of nuclei and scaled mean expression levels of cell-type classification markers corresponding to clusters shown in (**a**). **c**, Mean nuclei frequency of cell types across individual samples by genotype, separated into neuronal (top) and non-neuronal (bottom) populations. Data are presented as mean ± SEM and analyzed by mixed model test with Tukey”s multiple comparisons correction. \**p<*0.0332, \*\**p<*0.0021, \*\*\**p<*0.0002, and \*\*\*\**p<*0.0001. **d-f**, Quality control metrics per sample, including total nuclei (**d**), mean reads per nucleus (**e**), and mean number of unique genes detected per nucleus (**f**). **g**, Group-specific expression profiles projected onto the UMAP shown in (**a**). Notable transcriptomic shifts are observed within microglial clusters (closed arrows) and a subset of inhibitory neuron nuclei (open arrow). Data related to Fig. 3,5,6 and 7, and Extended Data Fig. 7–9. See Supplemental Tables 1 and 2 for cluster marker genes and cluster cell type classification. snRNAseq, single-nuclei RNA sequencing; UMAP, Uniform Manifold Approximation and Projection; Ex, excitatory; In Neuron, InN, inhibitory neuron; OPCs, oligodendrocyte precursor cells; OligoD, oligodendrocytes; CPEs, ciliary pigmented epithelial cells (ependymal cells); ABCs, arachnoid barrier cells; vLMCs, vascular leptomeningeal cells.

**Extended Data Fig. 7.**
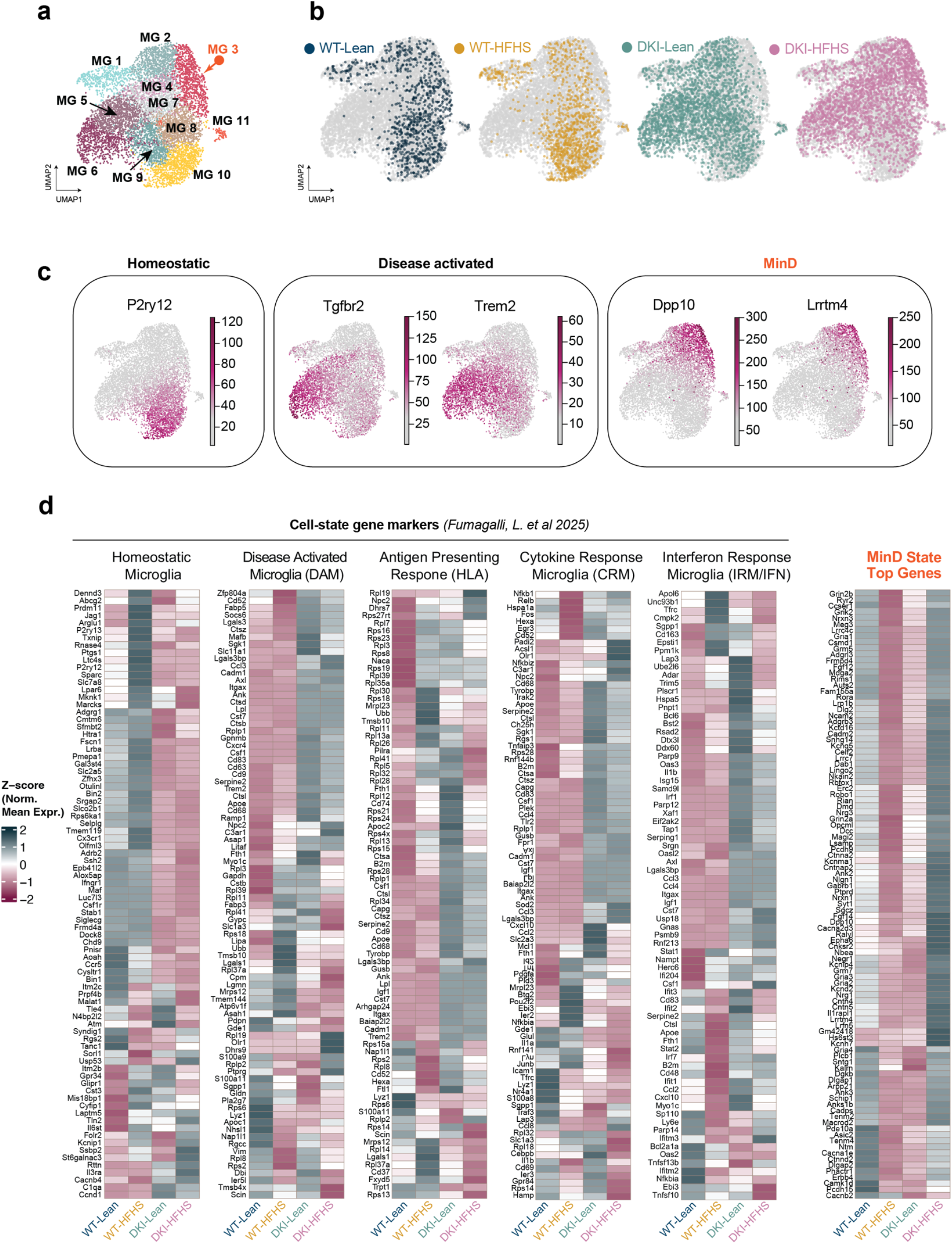
Disease-activated microglial signatures show HFHS-modulated expression in DKI mice. **a-b**, UMAP of microglia subclusters (**a**), and individual group expression profiles (**b**). Data related to Fig. 5a-c. **c**, Transcript levels of gene markers representing microglia homeostatic, disease activation, and MinD physiological states in UMAP space are shown in **a-b. d**, Comparative heatmaps showing the relative expression patterns of top genes representing curated microglia cell-state markers versus MinD state top genes.

**Extended Data Fig. 8.**
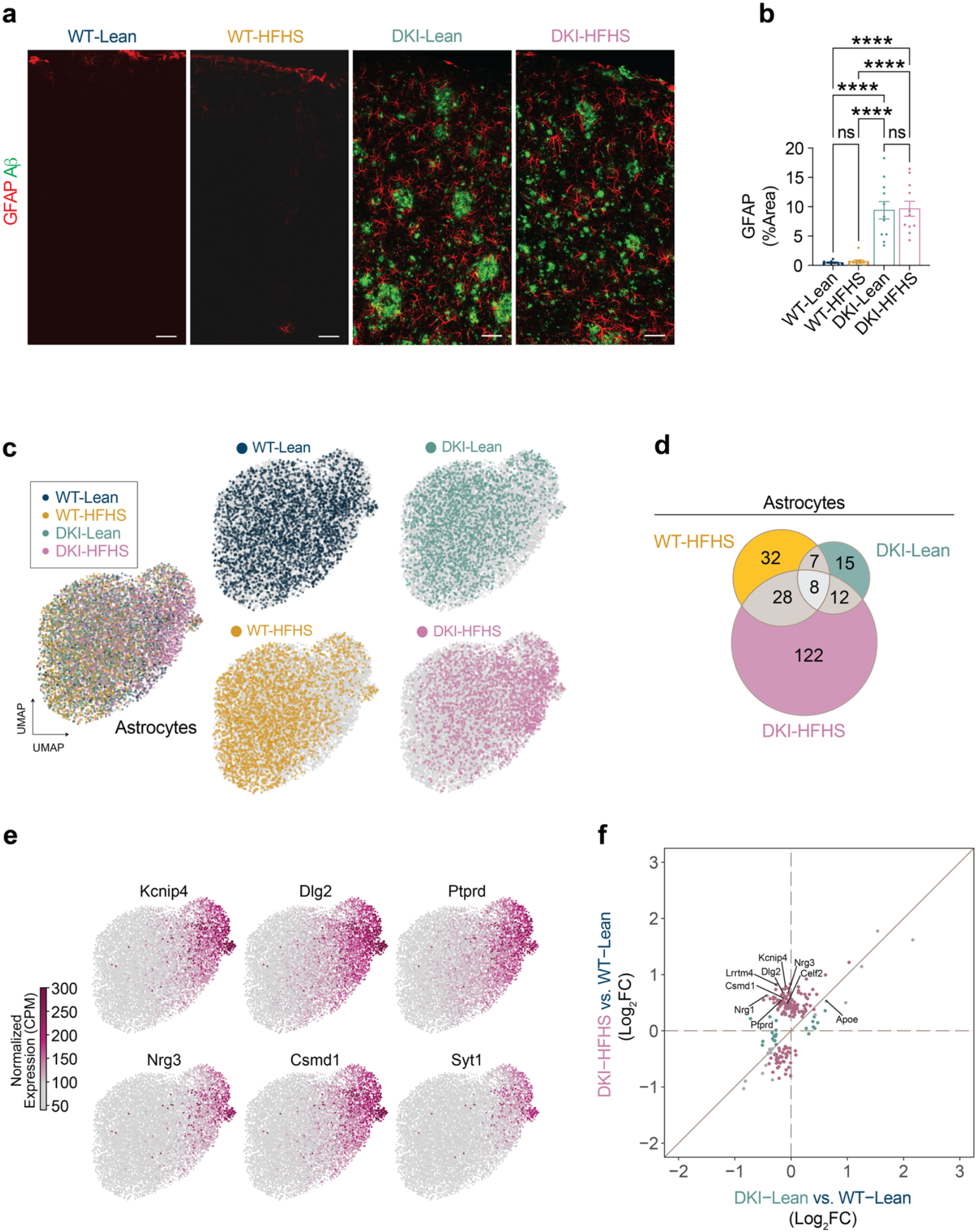
HFHS diet induces a MinD state in astrocyte transcriptomes of DKI mice. **a-f**, Transcriptomic assessment of astrocyte changes in response to diet, disease, and diet-disease interaction. **a**, Representative images of cortical astrocyte coverage (red) and amyloid-β deposition (green). **b**, Quantification of astrocyte coverage represented by GFAP signal in **a**. (Data are presented as mean ± SEM and analyzed by mixed model test with Tukey”s multiple comparisons correction. (For **b**, *n=9*, WT-Lean; *n=11*, for WT-HFHS, DKI- Lean, and DKI-HFHS groups. \**p<*0.0332, \*\**p<*0.0021, \*\*\**p<*0.0002, and \*\*\*\**p<*0.0001; ns: not significant) **c**, UMAP of astrocyte subcluster of overlayed (left) and individual (right) group expression profiles. **d**, Venn diagram showing shared and unique DEGs in astrocytes between WT-HFHS, DKI-Lean, and DKI-HFHS groups as compared to WT-Lean mice. **e**, Nuclei transcript levels of the top six marker genes for DKI-HFHS astrocytes. **f**, Correlation plot comparing Log_2_ fold-changes of DEGs in DKI-Lean and DKI-HFHS astrocytes relative to WT- Lean microglia. Date related to Fig. 4 and Extended Data Fig. 6. See Supplemental Table 4 for the complete DEG list. CPM: counts per million.

**Extended Data Fig. 9.**
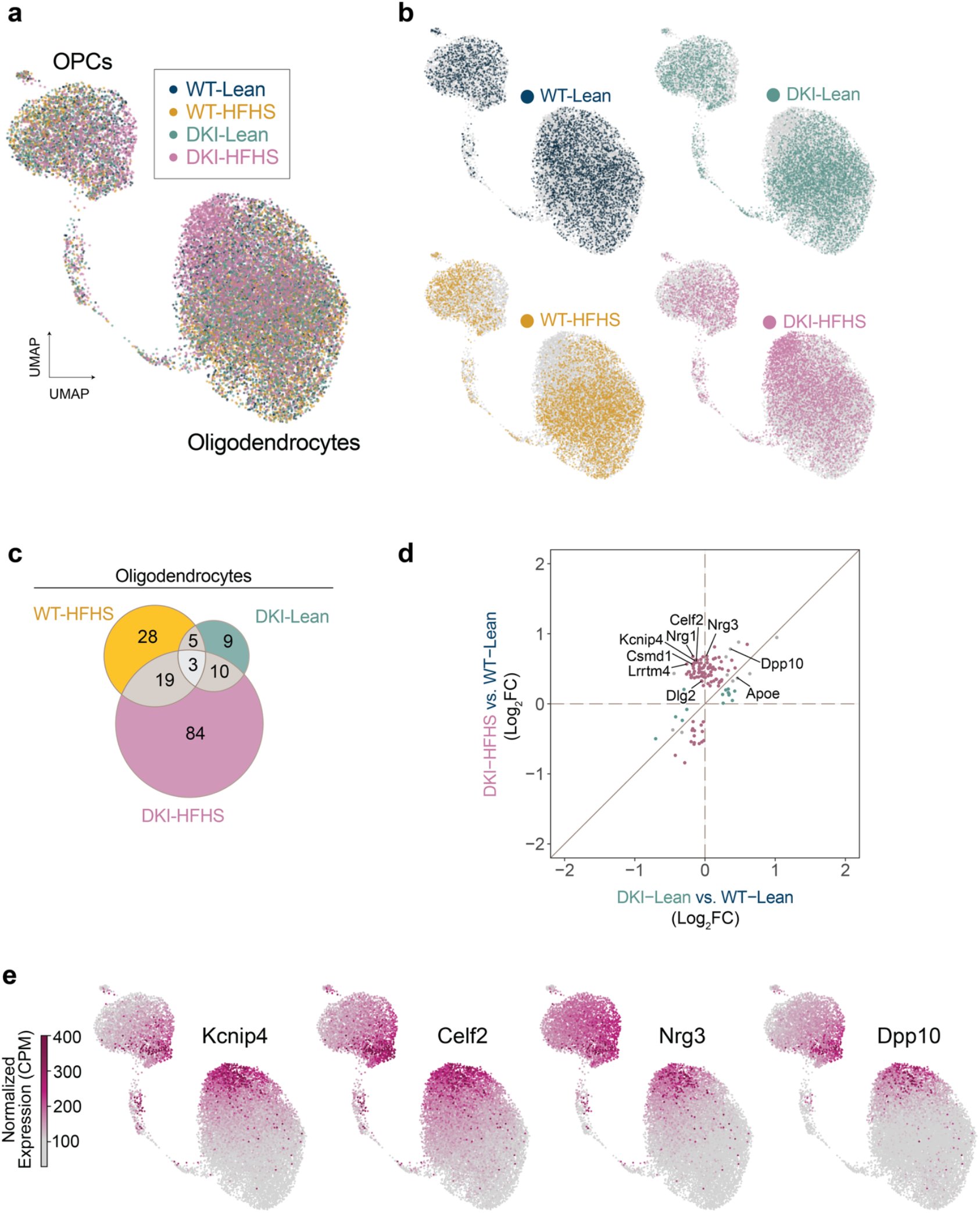
HFHS diet induces a MinD state in oligodendrocyte transcriptomes of DKI mice. **a-f**, Transcriptomics profiles of oligodendrocytes in response to diet, disease, and diet-disease interaction. **a-b**, UMAP of OPCs and oligodendrocytes subclusters of overlayed (**a**) and individual (**b**) group expression profiles. **c**, Venn diagram showing shared and unique differentially expressed genes (DEGs) in oligodendrocytes between WT-HFHS, DKI-Lean, and DKI-HFHS groups as compared to WT. **d**, Correlation plot comparing Log_2_ fold-changes of DEGs in DKI-Lean and DKI-HFHS astrocytes relative to WT-Lean microglia. **e**, Nuclei transcript levels of the top six marker genes for DKI-HFHS oligodendrocytes. Date related to Fig. 4 and Extended Data Fig. 6. See Supplemental Table 4 for the complete DEG list. ns: not significant; CPM: counts per million.

**Extended Data Fig. 10.**
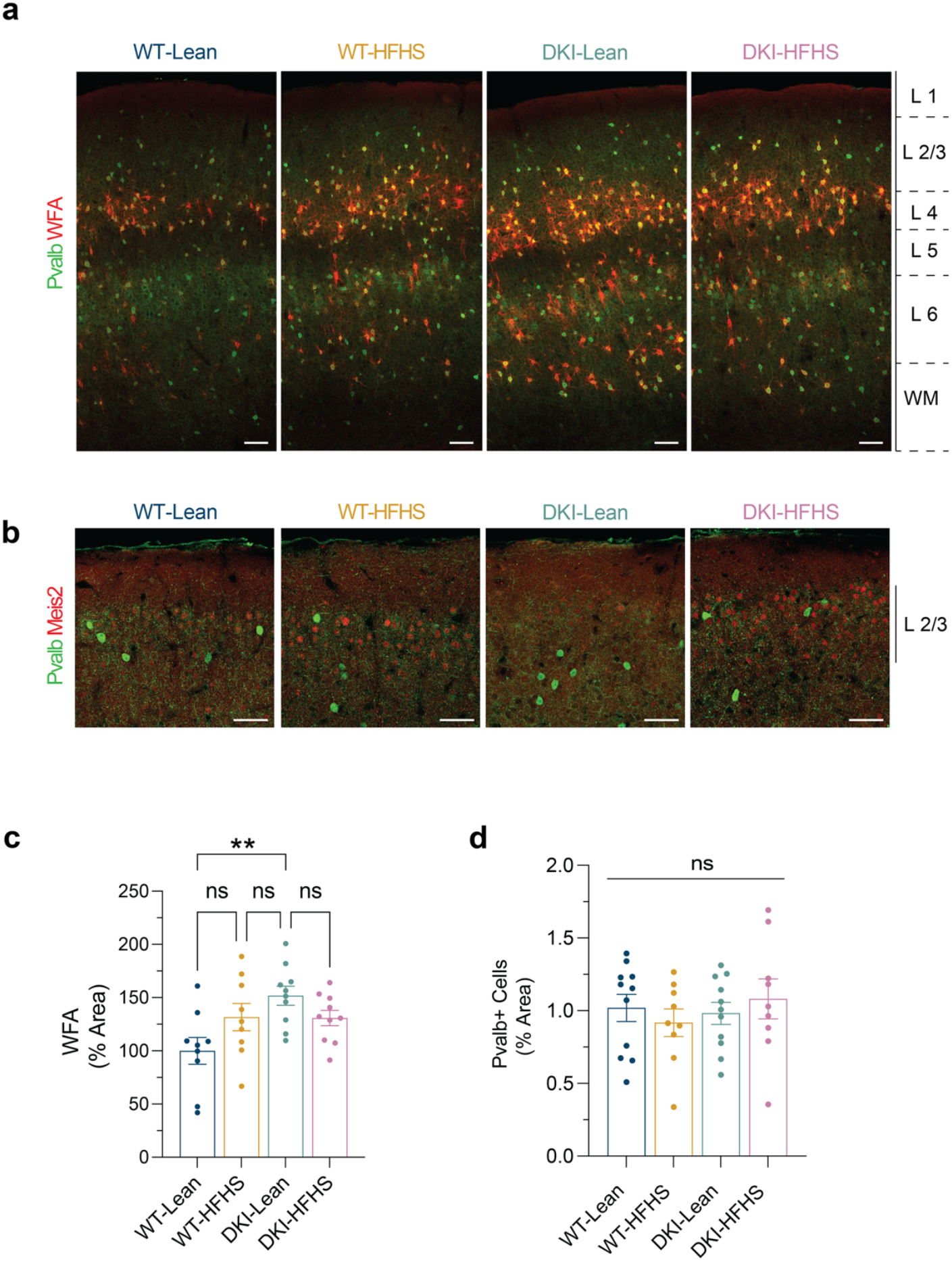
HFHS and disease induce changes in perineuronal nets in WT and DKI mice. **a**, Representative images showing WFA (red) and anti-Pvalb (Green) staining. Scale bars, 100 μm. **b**,**c**, Quantification of the percent area of WFA coverage (**b**) and Pvalb^+^ cells (**c**). (*n=9* for WT-Lean and WT-HFHS groups; *n=10* for DKI-Lean and DKI-HFHS groups). Data presented as the mean ± SEM and analyzed by two-way ANOVA with Tukey”s multiple comparisons test, \**p<*0.0332, \*\**p<*0.0021, \*\*\**p<*0.0002, and \*\*\*\**p<*0.0001; ns: not significant.

**Extended Data Fig. 11.**
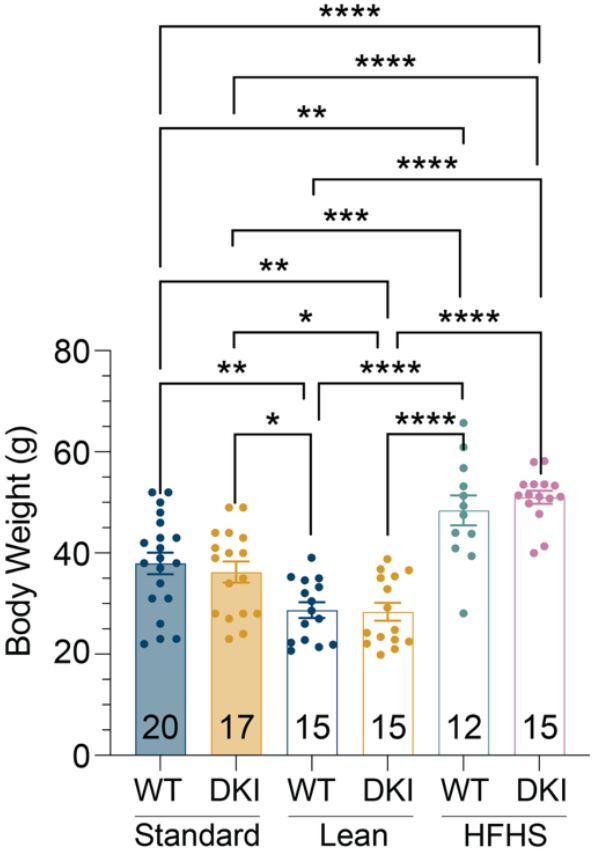
Body weight corresponds to diet nutrient composition. Body weight measurements were taken at 11.5 months of age in WT and DKI mice on Standard, Lean, or HFHS diet. The number of mice per group is displayed within each bar. Standard diet mice were fed *ad libitum* at the time of weaning. Mice on Lean or HFHS diets were fed *ad libitum* as described in the methods.

## Supplemental information

**Supplementary Table 1.** Gene expression marker list. Marker genes were determined using *Scanpy”s rank_gene_groups* function, applying the Wilcoxon rank-sum test, with adjustment for multiple comparisons via the Benjamini-Hochberg method.

**Supplementary Table 2.** Cell-type classifications as determined by the Allen Brain Atlas MapMyCells tool (RRID:SCR_024672).

**Supplementary Table 3.** Differentially expressed genes and pathway enrichment analysis for microglia.

**Supplementary Table 4.** Differentially expressed genes in astrocytes and oligodendrocytes.

**Supplementary Table 5.** Differentially expressed genes and pathway enrichment analysis for Meis2-positive inhibitory neurons.

**Supplementary Table 6.** Differentially expressed genes and pathway enrichment analysis for ExN 10 subcluster of excitatory neuron nuclei representing cortical layers 2-4.

## Notes

### Competing Interest Statement

The authors have declared no competing interest.

## References

1 Crane, P. K. et al. Glucose levels and risk of dementia. N Engl J Med 369, 540–548 (2013). 10.1056/NEJMoa1215740

2 Sun, Y. et al. Metabolism: A Novel Shared Link between Diabetes Mellitus and Alzheimer”s Disease. J Diabetes Res 2020, 4981814 (2020). 10.1155/2020/4981814

3 van Gils, V. et al. The association of glucose metabolism measures and diabetes status with Alzheimer”s disease biomarkers of amyloid and tau: A systematic review and meta-analysis. Neurosci Biobehav Rev 159, 105604 (2024). 10.1016/j.neubiorev.2024.105604

4 Lo, C. M. et al. Impaired insulin secretion and enhanced insulin sensitivity in cholecystokinin-deficient mice. Diabetes 60, 2000–2007 (2011). 10.2337/db10-0789

5 Liu, Z. et al. High-fat diet induces hepatic insulin resistance and impairment of synaptic plasticity. PLoS One 10, e0128274 (2015). 10.1371/journal.pone.0128274

6 Leclerc, M. et al. Cerebrovascular insulin receptors are defective in Alzheimer”s disease. Brain 146, 75–90 (2023). 10.1093/brain/awac309

7 Alonge, K. M., Porte, D., Jr. & Schwartz, M. W. Distinct Roles for Brain and Pancreas in Basal and Postprandial Glucose Homeostasis. Diabetes 72, 547–556 (2023). 10.2337/db22-0969

8 Barthold, D. et al. Alzheimer”s Disease-Related Neuropathology Among Patients with Medication Treated Type 2 Diabetes in a Community-Based Autopsy Cohort. J Alzheimers Dis 83, 1303–1312 (2021). 10.3233/JAD-210059

9 Peng, Y. et al. True or false? Alzheimer”s disease is type 3 diabetes: Evidences from bench to bedside. Ageing Res Rev 99, 102383 (2024). 10.1016/j.arr.2024.102383

10 Toledano, A., Rodriguez-Casado, A., Alvarez, M. I. & Toledano-Diaz, A. Alzheimer”s Disease, Obesity, and Type 2 Diabetes: Focus on Common Neuroglial Dysfunctions (Critical Review and New Data on Human Brain and Models). Brain Sci 14 (2024). 10.3390/brainsci14111101

11 Martemucci, G. et al. Metabolic Syndrome: A Narrative Review from the Oxidative Stress to the Management of Related Diseases. Antioxidants (Basel) 12 (2023). 10.3390/antiox12122091

12 Messiah, S. E. et al. Prevalence of the metabolic syndrome by household food insecurity status in the United States adolescent population, 2001-2020: a cross-sectional study. Am J Clin Nutr 119, 354–361 (2024). 10.1016/j.ajcnut.2023.11.014

13 Matsuzaki, T. et al. Insulin resistance is associated with the pathology of Alzheimer disease: the Hisayama study. Neurology 75, 764–770 (2010). 10.1212/WNL.0b013e3181eee25f

14 Schrijvers, E. M. et al. Insulin metabolism and the risk of Alzheimer disease: the Rotterdam Study. Neurology 75, 1982–1987 (2010). 10.1212/WNL.0b013e3181ffe4f6

15 Kim, A. B. & Arvanitakis, Z. Insulin resistance, cognition, and Alzheimer disease. Obesity (Silver Spring) 31, 1486–1498 (2023). 10.1002/oby.23761

16 Sai, K. K. S. et al. First-in-human positron emission tomography study of intranasal insulin in aging and MCI. Alzheimer”s & Dementia: Translational Research & Clinical Interventions 11, e70123 (2025). 10.1002/trc2.70123

17 Wang, W. et al. Associations of semaglutide with first-time diagnosis of Alzheimer”s disease in patients with type 2 diabetes: Target trial emulation using nationwide real-world data in the US. Alzheimer”s & Dementia 20, 8661–8672 (2024). 10.1002/alz.14313

18 de la Garza-Rodea, A. S., Knaäen-Shanzer, S., den Hartigh, J. D., Verhaegen, A. P. L. & van Bekkum, D. W. Anomer-equilibrated streptozotocin solution for the induction of experimental diabetes in mice (Mus musculus). J Am Assoc Lab Anim 49, 40–44 (2010).

19 Furman, B. L. Streptozotocin-Induced Diabetic Models in Mice and Rats. Curr Protoc 1, e78 (2021). 10.1002/cpz1.78

20 Winzell, M. S. & Ahren, B. The high-fat diet-fed mouse: a model for studying mechanisms and treatment of impaired glucose tolerance and type 2 diabetes. Diabetes 53 Suppl 3, S215–219 (2004). 10.2337/diabetes.53.suppl_3.s215

21 Andrikopoulos, S., Blair, A. R., Deluca, N., Fam, B. C. & Proietto, J. Evaluating the glucose tolerance test in mice. Am J Physiol Endocrinol Metab 295, E1323–1332 (2008). 10.1152/ajpendo.90617.2008

22 Saito, T. et al. Single App knock-in mouse models of Alzheimer”s disease. Nat Neurosci 17, 661–663 (2014). 10.1038/nn.3697

23 Chiasseu, M., Fesharaki-Zadeh, A., Saito, T., Saido, T. C. & Strittmatter, S. M. Gene-environment interaction promotes Alzheimer”s risk as revealed by synergy of repeated mild traumatic brain injury and mouse knock-in. Neurobiol Dis 145 (2020). 10.1016/j.nbd.2020.105059

24 Saito, T. et al. Humanization of the entire murine Mapt gene provides a murine model of pathological human tau propagation. J Biol Chem 294, 12754–12765 (2019). 10.1074/jbc.RA119.009487

25 Spurrier, J. et al. Reversal of synapse loss in Alzheimer mouse models by targeting mGluR5 to prevent synaptic tagging by C1Q. Sci Transl Med 14, eabi8593 (2022). 10.1126/scitranslmed.abi8593

26 Stoner, A. et al. Neuronal transcriptome, tau and synapse loss in Alzheimer”s knock-in mice require prion protein. Alzheimers Res Ther 15, 201 (2023). 10.1186/s13195-023-01345-z

27 Amelianchik, A., Sweetland-Martin, L. & Norris, E. H. The effect of dietary fat consumption on Alzheimer”s disease pathogenesis in mouse models. Translational Psychiatry 12, 293 (2022). 10.1038/s41398-022-02067-w

28 Valentin-Escalera, J., Leclerc, M., Calon, F. & Panza, F. High-Fat Diets in Animal Models of Alzheimer”s Disease: How Can Eating Too Much Fat Increase Alzheimer”s Disease Risk? Journal of Alzheimer”s Disease 97, 977–1005 (2024). 10.3233/jad-230118

29 Omar, B., Pacini, G. & Ahren, B. Differential development of glucose intolerance and pancreatic islet adaptation in multiple diet induced obesity models. Nutrients 4, 1367–1381 (2012). 10.3390/nu4101367

30 Ayala, J. E. et al. Standard operating procedures for describing and performing metabolic tests of glucose homeostasis in mice. Dis Model Mech 3, 525–534 (2010). 10.1242/dmm.006239

31 Spurrier, J. et al. Reversal of synapse loss in Alzheimer mouse models by targeting mGluR5 to prevent synaptic tagging by C1Q. Science Translational Medicine 14 (2022). 10.1126/scitranslmed.abi8593

32 Keren-Shaul, H. et al. A Unique Microglia Type Associated with Restricting Development of Alzheimer”s Disease. Cell 169, 1276–1290 e1217 (2017). 10.1016/j.cell.2017.05.018

33 Fumagalli, L. et al. Microglia heterogeneity, modeling and cell-state annotation in development and neurodegeneration. Nat Neurosci (2025). 10.1038/s41593-025-01931-4

34 Krasemann, S. et al. The TREM2-APOE Pathway Drives the Transcriptional Phenotype of Dysfunctional Microglia in Neurodegenerative Diseases. Immunity 47, 566–581 e569 (2017). 10.1016/j.immuni.2017.08.008

35 Kulkarni, B., Kumar, D., Cruz-Martins, N. & Sellamuthu, S. Role of TREM2 in Alzheimer”s Disease: A Long Road Ahead. Mol Neurobiol 58, 5239–5252 (2021). 10.1007/s12035-021-02477-9

36 Wang, S. et al. TREM2 drives microglia response to amyloid-β via SYK-dependent and -independent pathways. Cell 185, 4153–4169.e4119 (2022). 10.1016/j.cell.2022.09.033

37 Xiang, X. et al. TREM2 deficiency reduces the efficacy of immunotherapeutic amyloid clearance. EMBO Mol Med 8, 992–1004 (2016). 10.15252/emmm.201606370

38 Qiao, W. et al. Trem2 H157Y increases soluble TREM2 production and reduces amyloid pathology. Mol Neurodegener 18, 8 (2023). 10.1186/s13024-023-00599-3

39 Lee, S.-H. et al. Trem2 restrains the enhancement of tau accumulation and neurodegeneration by β-amyloid pathology. Neuron 109, 1283–1301.e1286 (2021). 10.1016/j.neuron.2021.02.010

40 Masuda, T. et al. IRF8 Is a Critical Transcription Factor for Transforming Microglia into a Reactive Phenotype. Cell Reports 1, 334–340 (2012). 10.1016/j.celrep.2012.02.014

41 Saeki, K., Pan, R., Lee, E., Kurotaki, D. & Ozato, K. IRF8 defines the epigenetic landscape in postnatal microglia, thereby directing their transcriptome programs. Nature Immunology 25, 1928–1942 (2024). 10.1038/s41590-024-01962-2

42 Ledonne, A. & Mercuri, N. B. On the Modulatory Roles of Neuregulins/ErbB Signaling on Synaptic Plasticity. Int J Mol Sci 21 (2019). 10.3390/ijms21010275

43 Liu, J. et al. Prevention of Alzheimer Pathology by Blocking Neuregulin Signaling on Microglia. eNeuro 10 (2023). 10.1523/ENEURO.0422-23.2023

44 Woo, R. S. et al. Neuregulin-1 enhances depolarization-induced GABA release. Neuron 54, 599–610 (2007). 10.1016/j.neuron.2007.04.009

45 Batista-Brito, R. et al. Developmental loss of ErbB4 in PV interneurons disrupts state-dependent cortical circuit dynamics. Mol Psychiatry 28, 3133–3143 (2023). 10.1038/s41380-023-02066-3

46 Spahic, H. et al. Dysregulation of ErbB4 Signaling Pathway in the Dorsal Hippocampus after Neonatal Hypoxia-Ischemia and Late Deficits in PV(+) Interneurons, Synaptic Plasticity and Working Memory. Int J Mol Sci 24 (2022). 10.3390/ijms24010508

47 Segerstolpe, Å. et al. Single-Cell Transcriptome Profiling of Human Pancreatic Islets in Health and Type 2 Diabetes. Cell Metabolism 24, 593–607 (2016). 10.1016/j.cmet.2016.08.020

48 Ottosson-Laakso, E. et al. Glucose-Induced Changes in Gene Expression in Human Pancreatic Islets: Causes or Consequences of Chronic Hyperglycemia. Diabetes 66, 3013–3028 (2017). 10.2337/db17-0311

49 Beddows, C. A. et al. Pathogenic hypothalamic extracellular matrix promotes metabolic disease. Nature 633, 914–922 (2024). 10.1038/s41586-024-07922-y

50 Fawcett, J. W. et al. The extracellular matrix and perineuronal nets in memory. Mol Psychiatry 27, 3192–3203 (2022). 10.1038/s41380-022-01634-3

51 Lupori, L. et al. A comprehensive atlas of perineuronal net distribution and colocalization with parvalbumin in the adult mouse brain. Cell Rep 42, 112788 (2023). 10.1016/j.celrep.2023.112788

52 Dong, H. et al. Metabolic memory: mechanisms and diseases. Signal Transduct Target Ther 9, 38 (2024). 10.1038/s41392-024-01755-x

53 Xiang, Q. et al. Insulin resistance-induced hyperglycemia decreased the activation of Akt/CREB in hippocampus neurons: Molecular evidence for mechanism of diabetes-induced cognitive dysfunction. Neuropeptides 54, 9–15 (2015). 10.1016/j.npep.2015.08.009

54 Arvanitakis, Z. et al. Brain Insulin Signaling, Alzheimer Disease Pathology, and Cognitive Function. Ann Neurol 88, 513–525 (2020). 10.1002/ana.25826

55 Gurley, S. B. et al. Impact of genetic background on nephropathy in diabetic mice. Am J Physiol Renal Physiol 290, F214–222 (2006). 10.1152/ajprenal.00204.2005

56 Takeda, S. et al. Diabetes-accelerated memory dysfunction via cerebrovascular inflammation and Aβ deposition in an Alzheimer mouse model with diabetes. Proceedings of the National Academy of Sciences 107, 7036–7041 (2010). 10.1073/pnas.1000645107

57 Wang, W. et al. Effects of high-fat diet on nutrient metabolism and cognitive functions in young APPKINL-G-F/NL-G-F mice. Neuropsychopharm Rep 42, 272–280 (2022). 10.1002/npr2.12257

58 Garcia-Serrano, A. M. & Duarte, J. M. N. Brain Metabolism Alterations in Type 2 Diabetes: What Did We Learn From Diet-Induced Diabetes Models? Front Neurosci 14, 229 (2020). 10.3389/fnins.2020.00229

59 Chen, X. et al. Microglia-mediated T cell infiltration drives neurodegeneration in tauopathy. Nature 615, 668–677 (2023). 10.1038/s41586-023-05788-0

60 Escoubas, C. C. et al. Type-I-interferon-responsive microglia shape cortical development and behavior. Cell 187, 1936–1954 e1924 (2024). 10.1016/j.cell.2024.02.020

61 Yanaizu, M., Washizu, C., Nukina, N., Satoh, J. I. & Kino, Y. CELF2 regulates the species-specific alternative splicing of TREM2. Sci Rep 10, 17995 (2020). 10.1038/s41598-020-75057-x

62 Shaw, B. C. et al. An Alternatively Spliced TREM2 Isoform Lacking the Ligand Binding Domain is Expressed in Human Brain. J Alzheimers Dis 87, 1647–1657 (2022). 10.3233/JAD-215602

63 Iguchi, A. et al. INPP5D modulates TREM2 loss-of-function phenotypes in a beta-amyloidosis mouse model. iScience 26, 106375 (2023). 10.1016/j.isci.2023.106375

64 Mei, L. & Nave, K. A. Neuregulin-ERBB signaling in the nervous system and neuropsychiatric diseases. Neuron 83, 27–49 (2014). 10.1016/j.neuron.2014.06.007

65 Luo, B. et al. ErbB4 promotes inhibitory synapse formation by cell adhesion, independent of its kinase activity. Transl Psychiatry 11, 361 (2021). 10.1038/s41398-021-01485-6

66 Santiago-Marrero, I. et al. Energy Expenditure Homeostasis Requires ErbB4, an Obesity Risk Gene, in the Paraventricular Nucleus. eNeuro 10 (2023). 10.1523/ENEURO.0139-23.2023

67 Ji, G., Li, S., Ye, L. & Guan, J. Gene Module Analysis Reveals Cell-Type Specificity and Potential Target Genes in Autism”s Pathogenesis. Biomedicines 9 (2021). 10.3390/biomedicines9040410

68 Pass, R. et al. Selective behavioural impairments in mice heterozygous for the cross disorder psychiatric risk gene DLG2. Genes Brain Behav 21, e12799 (2022). 10.1111/gbb.12799

69 Dhanasekara, C. S. et al. Association Between Autism Spectrum Disorders and Cardiometabolic Diseases: A Systematic Review and Meta-analysis. JAMA Pediatr 177, 248–257 (2023). 10.1001/jamapediatrics.2022.5629

70 Swift, G. H. et al. An endocrine-exocrine switch in the activity of the pancreatic homeodomain protein PDX1 through formation of a trimeric complex with PBX1b and MRG1 (MEIS2). Mol Cell Biol 18, 5109–5120 (1998). 10.1128/MCB.18.9.5109

71 Liu, Y., MacDonald, R. J. & Swift, G. H. DNA binding and transcriptional activation by a PDX1.PBX1b.MEIS2b trimer and cooperation with a pancreas-specific basic helix-loop-helix complex. J Biol Chem 276, 17985–17993 (2001). 10.1074/jbc.M100678200

72 Zhang, X. et al. Pax6 is regulated by Meis and Pbx homeoproteins during pancreatic development. Developmental Biology 300, 748–757 (2006). 10.1016/j.ydbio.2006.06.030

73 Krasnytska, D. O. et al. ERN1 dependent impact of glucose and glutamine deprivations on PBX3, PBXIP1, PAX6, MEIS1, and MEIS2 genes expression in U87 glioma cells. Endocr Regul 57, 37–47 (2023). 10.2478/enr-2023-0005

74 Biel, A. et al. AUTS2 Syndrome: Molecular Mechanisms and Model Systems. Front Mol Neurosci 15, 858582 (2022). 10.3389/fnmol.2022.858582

75 Pan, C. et al. The Kv2.2 channel mediates the inhibition of prostaglandin E2 on glucose-stimulated insulin secretion in pancreatic beta-cells. Elife 13 (2025). 10.7554/eLife.97234

76 Benner, O., Karr, C. H., Quintero-Gonzalez, A., Tamkun, M. M. & Chanda, S. The Shab family potassium channels are highly enriched at the presynaptic terminals of human neurons. J Biol Chem 301, 108235 (2025). 10.1016/j.jbc.2025.108235

77 Li, Z. et al. Protein Kinase C Controls the Excitability of Cortical Pyramidal Neurons by Regulating Kv2.2 Channel Activity. Neurosci Bull 38, 135–148 (2022). 10.1007/s12264-021-00773-x

78 Arnold, S. E. et al. Brain insulin resistance in type 2 diabetes and Alzheimer disease: concepts and conundrums. Nat Rev Neurol 14, 168–181 (2018). 10.1038/nrneurol.2017.185

79 Craft, S. et al. Safety, Efficacy, and Feasibility of Intranasal Insulin for the Treatment of Mild Cognitive Impairment and Alzheimer Disease Dementia: A Randomized Clinical Trial. JAMA Neurol 77, 1099–1109 (2020). 10.1001/jamaneurol.2020.1840

80 Alosco, M. L. et al. Modeling the Relationships Among Late-Life Body Mass Index, Cerebrovascular Disease, and Alzheimer”s Disease Neuropathology in an Autopsy Sample of 1,421 Subjects from the National Alzheimer”s Coordinating Center Data Set. J Alzheimers Dis 57, 953–968 (2017). 10.3233/JAD-161205

81 Sun, Z. et al. Late-life obesity is a protective factor for prodromal Alzheimer”s disease: a longitudinal study. Aging (Albany NY) 12, 2005–2017 (2020). 10.18632/aging.102738

82 Like, A. A. & Rossini, A. A. Streptozotocin-induced pancreatic insulitis: new model of diabetes mellitus. Science 193, 415–417 (1976). 10.1126/science.180605

83 de la Garza-Rodea, A. S., Knaan-Shanzer, S., den Hartigh, J. D., Verhaegen, A. P. & van Bekkum, D. W. Anomer-equilibrated streptozotocin solution for the induction of experimental diabetes in mice (Mus musculus). J Am Assoc Lab Anim Sci 49, 40–44 (2010).

84 Wolf, F. A., Angerer, P. & Theis, F. J. SCANPY: large-scale single-cell gene expression data analysis. Genome Biol 19, 15 (2018). 10.1186/s13059-017-1382-0

85 Traag, V. A., Waltman, L. & van Eck, N. J. From Louvain to Leiden: guaranteeing well-connected communities. Sci Rep 9, 5233 (2019). 10.1038/s41598-019-41695-z

86 Gayoso, A. et al. A Python library for probabilistic analysis of single-cell omics data. Nat Biotechnol 40, 163–166 (2022). 10.1038/s41587-021-01206-w

87 Shannon, P. et al. Cytoscape: a software environment for integrated models of biomolecular interaction networks. Genome Res 13, 2498–2504 (2003). 10.1101/gr.1239303

88 Bindea, G. et al. ClueGO: a Cytoscape plug-in to decipher functionally grouped gene ontology and pathway annotation networks. Bioinformatics 25, 1091–1093 (2009). 10.1093/bioinformatics/btp101

89 Szklarczyk, D. et al. The STRING database in 2021: customizable protein-protein networks, and functional characterization of user-uploaded gene/measurement sets. Nucleic Acids Res 49, D605–d612 (2021). 10.1093/nar/gkaa1074

